# Antibody responses in *Klebsiella pneumoniae* bloodstream infection: a cohort study

**DOI:** 10.1101/2024.05.01.591958

**Authors:** Wontae Hwang, Paeton L. Wantuch, Biana Bernshtein, Julia Zhiteneva, Damien Slater, Kian Hutt Vater, Sushmita Sridhar, Elizabeth Oliver, David J. Roach, Sowmya Rao, Sarah E. Turbett, Cory J. Knoot, Christian M. Harding, Mohammed Nurul Amin, Alan S. Cross, Regina C. LaRocque, David A. Rosen, Jason B. Harris

## Abstract

**Background:** *Klebsiella pneumonia* (Kpn) is the fourth leading cause of infection-related deaths globally, yet little is known about human antibody responses to invasive Kpn. In this study, we sought to determine whether the O-specific polysaccharide (OPS) antigen, a vaccine candidate, is immunogenic in humans with Kpn bloodstream infection (BSI). We also sought to define the cross-reactivity of human antibody responses among structurally related Kpn OPS subtypes and to assess the impact of capsule production on OPS-targeted antibody binding and function.

**Methods:** We measured plasma antibody responses to OPS (and MrkA, a fimbrial protein) in a cohort of patients with Kpn BSI and compared these with controls, including a cohort of healthy individuals and a cohort of individuals with *Enterococcus* BSI. We performed flow cytometry to measure the impact of Kpn capsule production on whole cell antibody binding and complement deposition, utilizing patient isolates with variable levels of capsule production and isogenic capsule-deficient strains derived from these isolates.

**Findings:** We enrolled 69 patients with Kpn BSI. Common OPS serotypes accounted for 57/69 (83%) of infections. OPS was highly immunogenic in patients with Kpn BSI, and peak OPS-IgG antibody responses in patients were 10 to 30-fold higher than antibody levels detected in healthy controls, depending on the serotype. There was significant cross-reactivity among structurally similar OPS subtypes, including the O1v1/O1v2, O2v1/O2v2 and O3/O3b subtypes. Physiological amounts of capsule produced by both hyperencapsulated and non-hyperencapsulated Kpn significantly inhibited OPS-targeted antibody binding and function.

**Interpretation:** OPS was highly immunogenic in patients with Kpn BSI, supporting its potential as a candidate vaccine antigen. The strong cross-reactivity observed between similar OPS subtypes in humans with Kpn BSI suggests that it may not be necessary to include all subtypes in an OPS-based vaccine. However, these observations are tempered by the fact that capsule production, even in non-highly encapsulated strains, has the potential to interfere with OPS antibody binding. This may limit the effectiveness of vaccines that exclusively target OPS.

**Funding:** National Institute of Allergy and Infectious Diseases at the National Institutes of Health.

**Research in Context:** *Evidence before this study:* Despite the potential of O-specific polysaccharide (OPS) as a vaccine antigen against *Klebsiella pneumoniae* (Kpn), the immunogenicity of OPS in humans remains largely unstudied, creating a significant knowledge gap with regard to vaccine development. A search of PubMed for publications up to March 18, 2024, using the terms “*Klebsiella pneumoniae*” and “O-specific polysaccharide” or “O-antigen” or “lipopolysaccharide” revealed no prior studies addressing OPS antibody responses in humans with Kpn bloodstream infections (BSI). One prior study^1^ evaluated antibody response to a single lipopolysaccharide (which contains one subtype of OPS) in humans with invasive Kpn infection; however, in this study OPS typing of the infecting strains and target antigen were not described.

*Added value of this study:* Our investigation into OPS immunogenicity in a human cohort marks a significant advance. Analyzing plasma antibody responses in 69 patients with Kpn BSI, we found OPS to be broadly immunogenic across all the types and subtypes examined, and there was significant cross-reactivity among structurally related OPS antigens. We also demonstrated that Kpn capsule production inhibit OPS antibody binding and the activation of complement on the bacterial surface, even in classical Kpn strains expressing lower levels of capsule.

*Implications of all the available evidence:* While the immunogenicity and broad cross-reactivity of OPS in humans with Kpn BSI suggests it is a promising vaccine candidate, the obstruction of OPS antibody binding and engagement by physiologic levels of Kpn capsule underscores the potential limitations of an exclusively OPS-antigen based vaccine for Kpn. Our study provides insights for the strategic development of vaccines aimed at combating Kpn infections, an important antimicrobial resistant pathogen.

## Introduction

*Klebsiella pneumoniae* (Kpn) is an increasing global concern, and ranks as the fourth leading cause of infection-related mortality, responsible for approximately 790,000 deaths annually.^2^ Kpn infections are caused by both classical (cKp) and hypervirulent (hvKp) pathotypes, which are distinguished by the ability of the latter to cause invasive infections in healthy individuals. Hypervirulent Kpn strains possess unique virulence factors, including siderophores and genes associated with capsule production.^3^ The global dissemination of hvKp, especially isolates that are resistant to antibiotics, poses a significant health threat.^4^ Despite the potential of vaccines against Kpn^5^, none are currently approved, and research is limited to pre-clinical stages, with only one vaccine trial registered in clinicaltrials.gov^6^and no Phase III trials yet.^7^

Given the need to develop vaccines for Kpn, there is an interest in identifying the antigenic repertoire of this pathogen and the associated correlates of protection^8^. Capsular polysaccharide (CPS) is one potential vaccine antigen since invasive Kpn are often encapsulated, and high levels of capsule expression may mask other bacterial cell surface targets, rendering non-CPS vaccines ineffective.^9^ However, not all invasive Kpn express abundant capsule, and in some infections CPS production may be downregulated.^10,11^ The diversity of the Kpn capsule also presents a challenge. Over 150 distinct Kpn CPS structures are predicted based on genomic analyses^12,13^, and it is estimated that a CPS-based vaccine would require the inclusion of more than 24 capsular antigens to achieve 60% coverage of the currently circulating invasive Kpn strains.^14^

The MrkA protein of Kpn, a key component of a type III fimbria, has also emerged as a target for vaccine development. MrkA’s conservation across strains and its role as an adhesin make it an attractive vaccine candidate, and studies have shown that immunization with MrkA can elicit strong IgG responses, significantly reducing bacterial burden and providing protection in animal models of Kpn, including models of sepsis and pneumonia.^15-17^

The O-specific polysaccharide (OPS), a component of lipopolysaccharide, is another target for Kpn vaccine development, due to its limited structural diversity.^12,14,18^ A recent survey estimated that population-based immunity to over 90% of invasive Kpn could be achieved with a quadrivalent O-polysaccharide antigen vaccine targeting the O1, O2, O3 and O5 serotypes.^14^ However, there are gaps in our understanding of the role of OPS antibody responses in immunity to Kpn. First, while OPS-based vaccines are protective in some mouse models of invasive Kpn,^9^ human antibody responses to OPS have not been characterized. Whether Kpn OPS is immunogenic in humans at risk for invasive Kpn is highly relevant to vaccine development. In addition, the extent to which heterologous OPS antigens induce cross-reactive antibody responses has implications for vaccine composition. Finally, it is unknown whether OPS induces a functional antibody response in humans, and whether OPS antibody binding is blocked by the variable levels of capsule produced by both hvKp and cKp.

To address these questions, we studied OPS antibody responses in patients with Kpn bloodstream infection (BSI). We measured the functional potential and cross-reactivity of OPS antibody responses in humans. We also determined whether physiologic amounts of capsule produced by invasive Kpn isolates interfered with OPS antibody binding and function. Our results demonstrate that OPS is a dominant antigen in humans with invasive Kpn, and that there is significant cross-reactivity among antibody responses to structurally similar OPS antigens. However, OPS targeted responses may be ineffective in inducing protection against Kpn, given that physiologic amounts of CPS production limited OPS antibody binding and function.

## Methods

### Study design and participants

Our investigation was carried out at Massachusetts General Hospital (MGH) a 1000-bed tertiary care hospital which provides care for patients of all ages. We enrolled all identified patients with Kpn BSI between 07/24/21 and 08/04/22. Demographic and clinical data, including data on immunosuppression, comorbidities, and other risk factors for invasive infection were extracted from the medical record, as described previously.^19^ Plasma was collected longitudinally for routine patient care over the course of each patients’ hospitalization or on subsequent follow-up visits. We excluded patients who had inadequate plasma collected due to low sample volume, death or lack of follow-up, as well as patients whose isolates were not confirmed to be *Klebsiella pneumoniae* by whole genome sequencing (WGS). hvKp were defined as strain which were *rmpA, iro*, and *iuc* positive.^20^ We compared immune responses in this Kpn BSI cohort with a previously described cohort of healthy adults (healthy controls, HC) presenting for a routine outpatient consultation at MGH^21^, and with a contemporaneously enrolled cohort of patients with *Enterococcus* BSI (BSI controls, BC) hospitalized at MGH.

### Procedures

Detailed procedures are included in the Supplemental Methods (appendix pp 3-9). In brief, plasma IgG, IgM and IgA antibody responses were measured using a customized multiplexed bead assay (MBA) to a panel of antigens including Exoprotein A of *Pseudomonas aeruginosa* (EPA) and Human Serum Albumin (HSA) conjugated OPS antigens, and MrkA. A plasma dilution series, prepared from a mixture of patient serum, was used to normalize the Median Fluorescence Intensity (MFI) values between experiments. Functional measures of the immune response included antibody-dependent complement deposition (ADCD), and antibody-dependent neutrophil phagocytosis (ADNP) targeting the O1v1 and O3b antigen. We measured ADNP responses using flow cytometry to detect the phagocytosis of antigen-coupled beads by donor-derived granulocytes, and ADCD responses using antigens conjugated with Luminex Magplex carboxylated beads incubated with guinea pig complement measured by flow cytometry as described previously.^22^ For selected patients, plasma antibody binding to whole bacterial cells was measured by flow cytometry using each patient’s paired plasma and Kpn isolate, which were engineered to constitutively express Green Fluorescent Protein (GFP), as well as plasma paired with an isogenic capsule deficient mutant derived from each patient’s GFP-expressing Kpn isolate. Isogenic capsule deficient mutants were constructed by deletion of the *wcaJ* gene using CRISPR-Cas9 and Red recombineering systems. Capsule expression for each strain and its isogenic *ΔwcaJ* mutant was measured using glucuronic acid quantification. OPS antibodies in plasma and purified immunoglobulin were blocked with an excess of each specified antigen (including O1v1-EPA, O3b-EPA, and EPA).

### Statistical analysis

Categorical values were analyzed using the Chi-squared test, while continuous variables, including the demographic features of the study participants, were assessed using a two-tailed T-test. Trends in antibody responses over time were assessed using LOWESS regression. The Mann-Whitney U test was used to determine the statistical significance of differences between independent groups. For analyzing results, except for those from multiplexed bead assays investigating antibody responses, ADCD, and ADNP, Welch’s t-test was utilized. We used Pearson correlation analysis to investigate correlative relationships. All statistical analyses were performed using Python (v3.9.7) and GraphPad Prism 10.

## Results

We enrolled 129 patients with suspected Kpn BSI over the 12-month study period (appendix p 10). Of the 129 patients, 43 were excluded because of death or insufficient follow-up, and 17 were excluded because WGS identified *K. variicola* or *K. quasipneumoniae* spp. rather than Kpn, leaving 69 participants eligible for measurement of OPS antibody responses. Among the participants, 3/69 (4%) were infected with hvKp. There were no significant differences in the age, sex, or immunocompromised status between the 69 patients who were included and the 60 patients who were excluded from the immunologic analysis (table 1). Participants ranged from 0 to 92 years of age, with a median age of 68 years, and 64% were male. The patients had a median Charlson Comorbidity Index of 5.25 and 35/69 (51%) were immunocompromised, mostly due to chemotherapy-induced neutropenia. Our study also included 36 healthy controls and 25 similarly medically complex, contemporaneously enrolled patients with *Enterococcus* BSI (table 1).

**Table 1.**
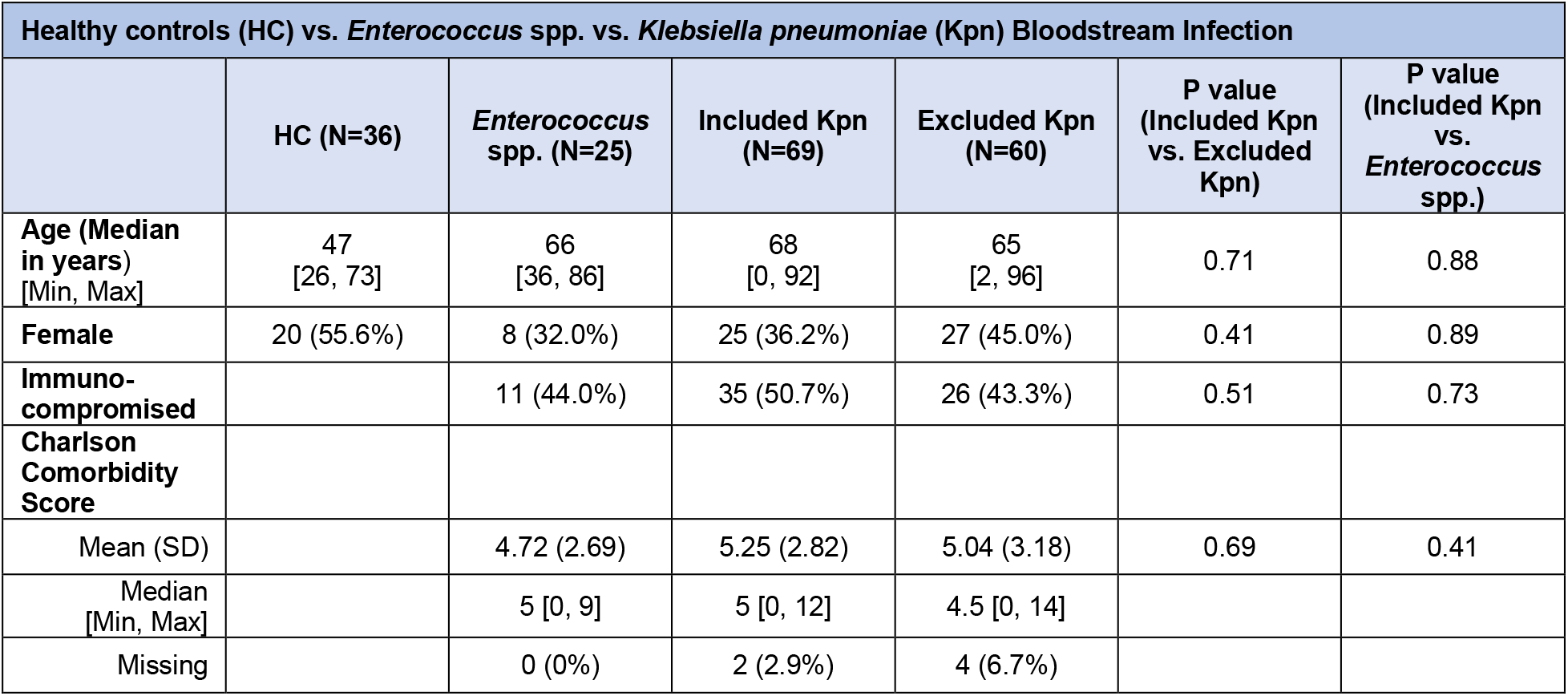
Demographic features of study participants.

| Healthy controls (HC) vs. *Enterococcus* spp. vs. *Klebsiella pneumoniae* (Kpn) Bloodstream Infection |  |  |  |  |  |  |
| --- | --- | --- | --- | --- | --- | --- |
|  | HC (N=36) | *Enterococcus* spp. (N=25) | Included Kpn (N=69) | Excluded Kpn (N=60) | P value (Included Kpn vs. Excluded Kpn) | P value (Included Kpn vs. *Enterococcus* spp.) |
| Age (Median in years) [Min, Max] | 47 [26, 73] | 66 [36, 86] | 68 [0, 92] | 65 [2, 96] | 0.71 | 0.88 |
| Female | 20 (55.6%) | 8 (32.0%) | 25 (36.2%) | 27 (45.0%) | 0.41 | 0.89 |
| Immuno-compromised |  | 11 (44.0%) | 35 (50.7%) | 26 (43.3%) | 0.51 | 0.73 |
| Charlson Comorbidity Score |  |  |  |  |  |  |
| Mean (SD) |  | 4.72 (2.69) | 5.25 (2.82) | 5.04 (3.18) | 0.69 | 0.41 |
| Median [Min, Max] |  | 5 [0, 9] | 5 [0, 12] | 4.5 [0, 14] |  |  |
| Missing |  | 0 (0%) | 2 (2.9%) | 4 (6.7%) |  |  |

Previous studies have shown that >80% of global Kpn infections are assigned to serotypes O1, O2, O3, or O5, which are considered potential OPS vaccine serotypes (figure 1A).^12,14^ O1 and O2 antigens contain repeating galactan subunits and O3 and O5 contain mannose repeats. In our cohort, these 4 serotypes accounted for 57/69 (83%) of all Kpn BSIs (figure 1B). The most identified CPS serotype was K2, which is historically associated with hvKp. However, no single CPS type predominated, and K2 was identified in only three patients (appendix pp 21-22).^4,23^

**Figure 1.**
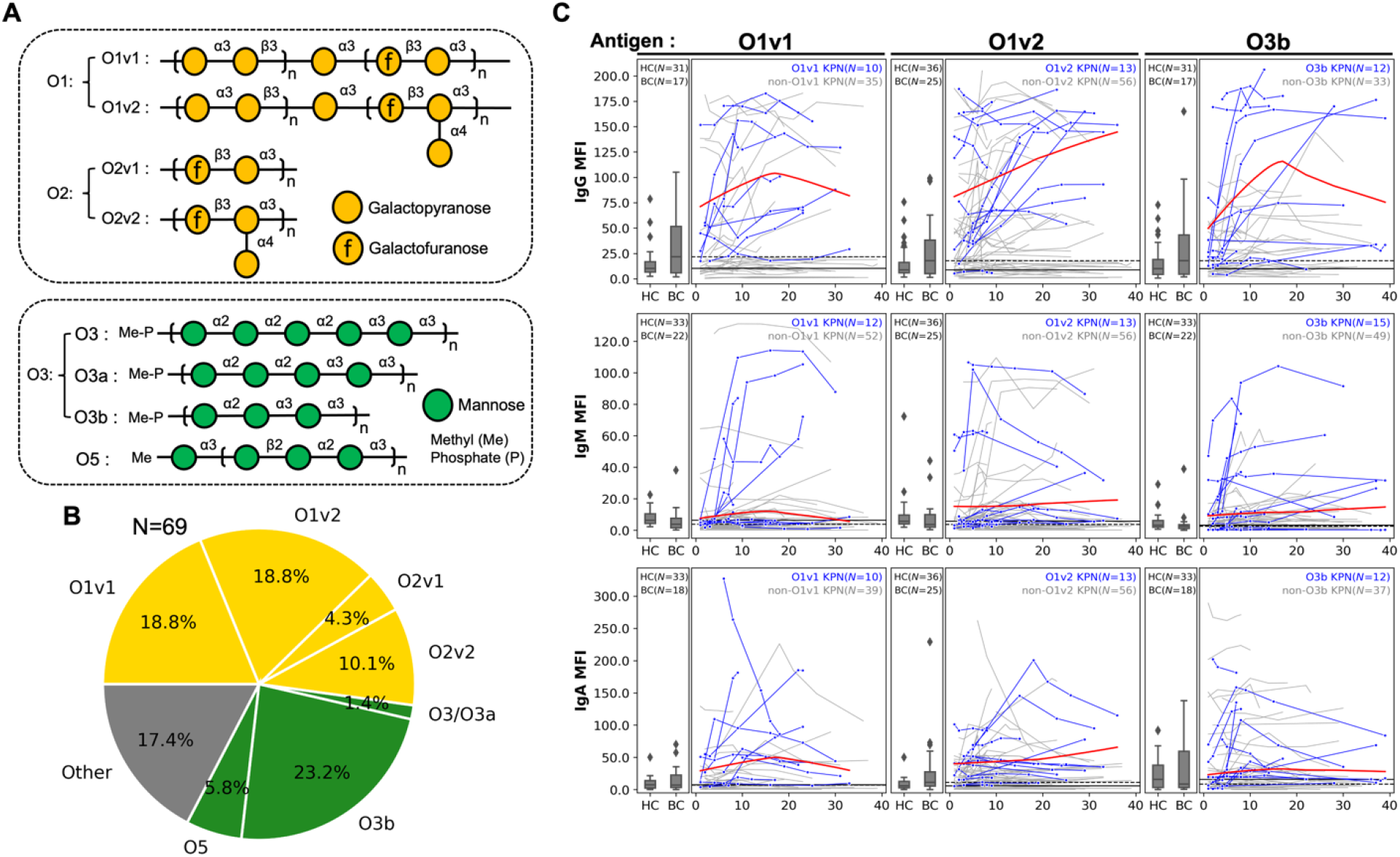
OPS serotypes and longitudinal antibody responses in patients with Kpn bloodstream infection (BSI) **(A)** The structures of O1, O2, O3, and O5 OPS are shown, with the galactan- (O1, O2) and mannan-based OPSs (O3, O5). **(B)** The results of *in silico* OPS serotyping for Kpn isolates. **(C)** IgG, IgM, and IgA antibody responses to O1v1, O1v2, and O3b antigens are shown. The Y-axis represents the median fluorescence intensity (MFI). Boxplots show median and interquartile MFI from healthy- (HC) and *Enterococcus* spp. bacteremic-controls (BC). Longitudinal antibody responses in patients with Kpn BSI are represented by individual lines, with each data point corresponding to a single sample at a given time point. The X-axis denotes the number of days since the positive blood culture for Kpn (Day 0). Blue lines indicate antibody responses of patients with Kpn BSI caused by infection the homologous OPS type, while gray lines represent patients Kpn BSI caused by heterologous OPS types. The red lines illustrate the LOWESS regression applied to the blue lines. The solid black line represents the median of HC, and the dashed black line represents the median of BC. The number of patients included is shown in each graph.

We measured plasma IgG, IgM, and IgA antibody responses to O1v1, O1v2, O2v1, O2v2, O3/O3a (O3), O3b, O5 and MrkA. Patients with Kpn BSI caused by O1v1, O1v2 and O3b strains, which were the most common in our cohort, exhibited elevated IgG, IgM and IgA antibody responses to their homologous OPS serotype relative to both healthy controls and *Enterococcus* BSI controls. In aggregate, OPS IgG responses increased over a 40-day period from the time of the initial Kpn positive blood culture (figure 1C and appendix p 11).

We then compared the peak OPS-IgG, IgM and IgA antibody responses of our cohort of patients with Kpn BSI to those in healthy controls and *Enterococcus* BSI controls (figure 2). Patients infected with O1v1, O1v2 and O3b Kpn demonstrated significant IgG, IgM and IgA responses to their homologous OPS compared to controls. Patients with BSI from the less common serotypes O2v1, O2v2, and O5 BSI demonstrated significant IgG or IgA responses compared to controls, but not all IgM responses achieved significance. Specifically, there was a 11.9-fold higher O1v1 IgG response (P=5.7E-06), a 17.8-fold higher O1v2 IgG response (P=1.2E-06), a 29.9-fold higher O2v1 IgG response (P=8.6E-04), a 12.0-fold higher O3b IgG response (P=1.2E-05), and a 16.8-fold higher O5 IgG response (P=0.032) in patients with Kpn BSI relative to healthy controls. Patients with Kpn BSI also had antibody responses to MrkA, with a 3.7-fold higher IgG response (P=7.2E-05) than healthy controls.

**Figure 2.**
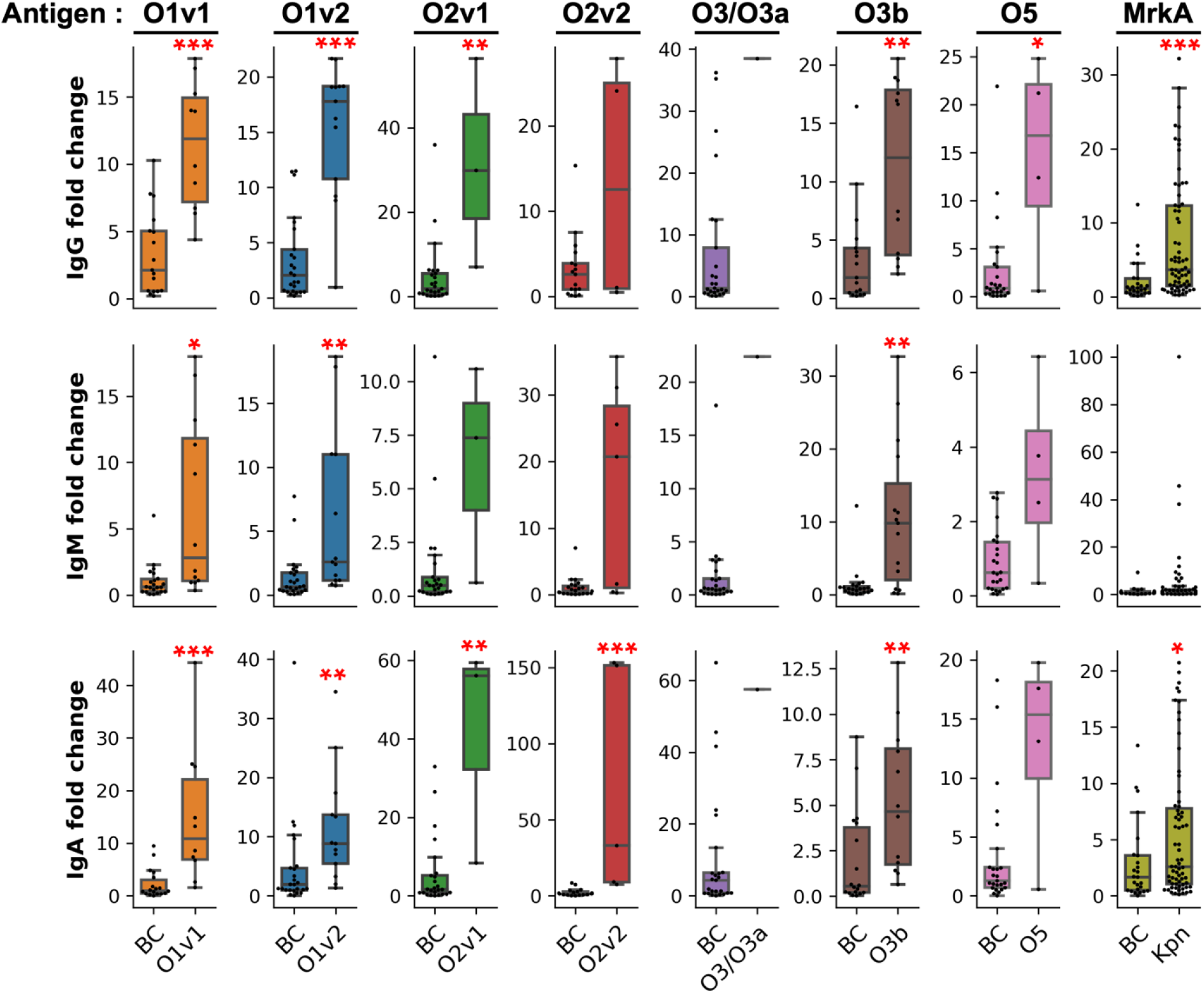
Elevated homologous OPS and MrkA antibody responses in Kpn BSI. Boxplots of the homologous OPS and MrkA antibody responses are shown in patients with Kpn BSI compared to patients with *Enterococcus* spp. BSI (BC). The Y-axis shows the MFI divided by the median MFI of healthy controls (HC). Asterisks indicate P values smaller than 0.05, as determined by Mann-Whitney U test, compared to both HC and BC. Significance levels are denoted as: * 0.01 ≤ P < 0.05, ** 0.001 ≤ P < 0.01, *** P < 0.001. The number of samples in each group is presented in the appendix (p 23).

We hypothesized that increased CPS expression might be associated with a lower antibody response to OPS, so we measured the correlation between CPS expression and the magnitude of the OPS antibody response. For this analysis, we focused on the O1v1 and O1v2 serotypes since these were the most common, and the best powered for this analysis. There was no correlation between the amount of CPS expressed by the infecting strain and the magnitude of the O1 antibody response in patients with Kpn O1 BSI (appendix p 12).

Many individuals at risk for invasive Kpn have compromised immunity, and our cohort included immunocompromised individuals. To assess the role of impaired immunity on the OPS antibody response, we compared OPS antibody responses in immunocompromised and non-immunocompromised patients infected with O1v1 or O1v2 Kpn. Both immunocompromised and immunocompetent patients had IgG and IgA antibody responses to OPS. In contrast, responses to MrkA were observed only in the immunocompetent patients (appendix pp 13-14). This suggests that OPS responses might be preserved in some immunocompromised individuals with Kpn BSI.

A cross-reactive response, which occurs when antigen stimulation generates antibodies which bind to structurally related antigens, can be immunologically advantageous. To assess OPS antibody cross-reactivity, we measured the correlation in antibody levels between different Kpn OPS subtypes (figure 3A). IgG, IgM and IgA antibody levels against O1v1 and O1v2 were all extremely correlated in the study participants (R^2^=0.96, P=7.1E-16 for IgG). Antibody levels were also correlated between the O2v1 and O2v2 subtypes (R^2^=0.61, P=4.8E-04 for IgG), and the O3/O3a and O3b subtypes (R^2^=0.70, P=2.4E-05 for IgG).

**Figure 3.**
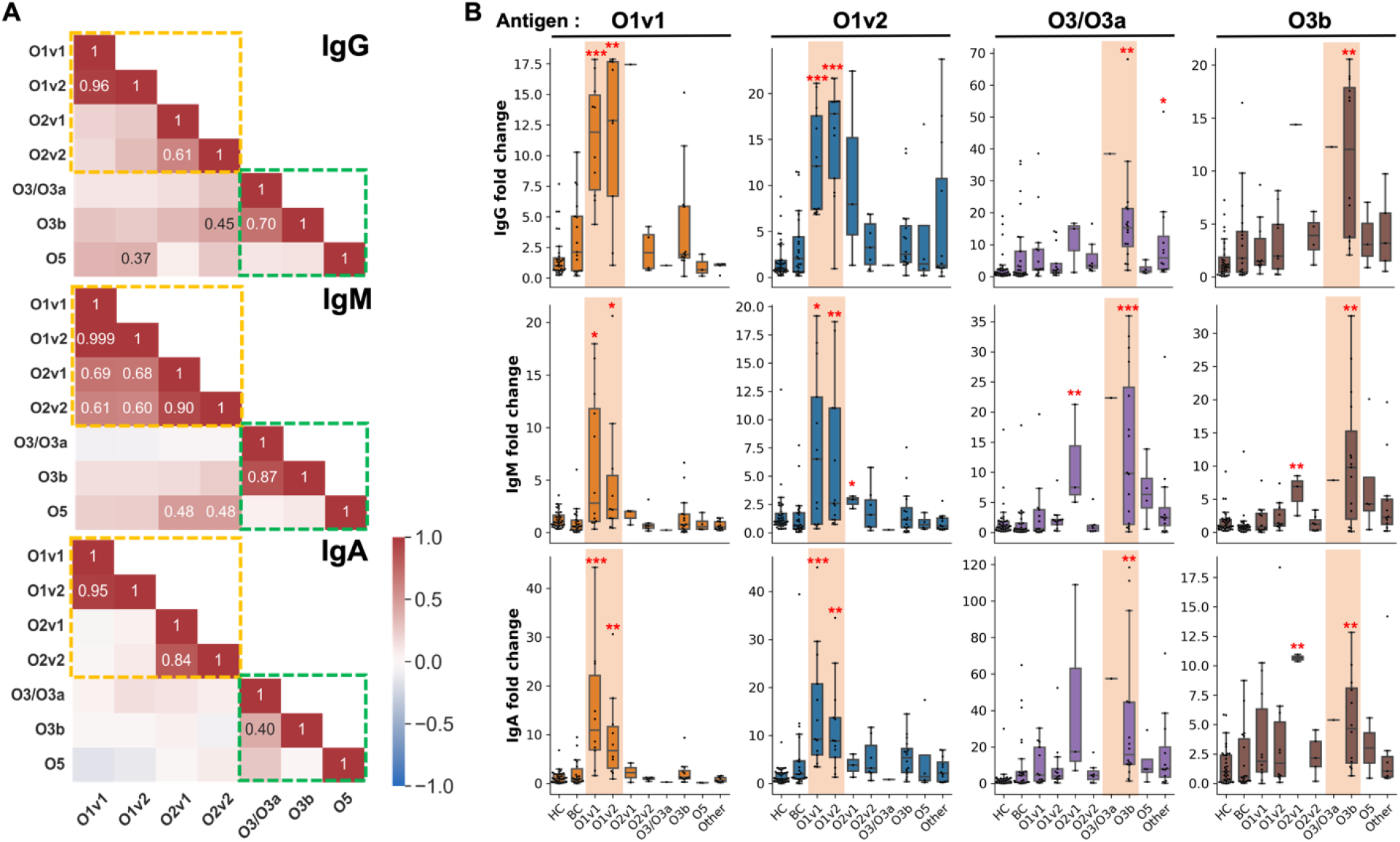
Cross-specific and cross-reactive antibody responses to closely related OPS. **(A)** A representative set of plasma samples with low cross-specificity to control antigens EPA and HSA were selected from patients with Kpn BSI, healthy controls (HC), and patients with *Enterococcus* spp. BSI (BC). The heatmap represents the correlation between MFIs for different OPS types, with numbers within the boxes indicating Pearson correlation values (R^2^) with a P value smaller than 0.05, while non-significant values are omitted. Galactan-based and mannan-based OPS results are distinguished by dashed orange and green lines. **(B)** Individual boxplots provide IgG, IgM, and IgA antibody responses for HC, BC, and the highest antibody responses observed in patients with Kpn BSI in response to O1v1, O1v2, O3/O3a (O3), and O3b antigens. The X-axis illustrates plasma samples sourced from HC, BC, and patients infected with Kpn presenting a diverse OPS type. The “Other” category encompasses OPS types O4, O12, and unidentified OPS. The Y-axis represents the fold change in antibody responses relative to HC, calculated as the MFI divided by the median MFI of HC. Specifically, the results from patients with Kpn BSI who exhibited closely related OPS, as described in (A), are highlighted with orange backgrounds. Red asterisks signify statistical significance (P values smaller than 0.05) when compared to both HC and BC using Mann-Whitney U test. Significance levels are denoted as follows: * for 0.01 ≤ P < 0.05, ** for 0.001 ≤ P < 0.01, and *** for P < 0.001. Detailed information regarding the number of samples is available in appendix (p 23).

To measure cross-reactive immune responses to OPS antigens, we next measured antibody responses to heterologous OPS antigens in patients with Kpn BSI (figure 3B). Patients with O1v1 or O1v2 Kpn BSI had a robust response to both the homologous and heterologous O1 subtypes. Patients with O3b Kpn BSI also had robust response to the heterologous O3/O3a (O3) subtype. Patients with O2v2 Kpn BSI also had a significant IgG and IgA response to the heterologous O2v1 antigen (e.g., 10.6-fold higher, P=0.001 for IgG compared to healthy controls). Due to the small numbers of patients with O2v1, O3 and O5 infection, we were unable to assess heterologous antibody responses following exposure to these antigens (appendix p 15).

To understand how OPS-targeted antibodies may mediate immune function, we measured OPS antibody-mediated ADNP and ADCD using OPS-conjugated beads. This analysis was conducted for patients infected with O1v1, O1v2, or O3b strains. Patients with O1v1 and O1v2 Kpn BSI had ADNP and ADCD responses to the O1v1 antigen, while patients with O3b infection had significant ADNP and ADCD responses to the homologous O3b antigen (appendix p 16).

After observing that Kpn BSI results in a functional OPS antibody response, we assessed whether capsule production might interfere with OPS antibody binding and ADCD responses. We evaluated antibody responses in four patients with either O1 or O3b Kpn BSI. The four patients were selected based on variable levels of CPS expression in their infecting strain, and included two patients infected with highly encapsulated Kpn (patient 24 was infected with hypermucoid cKp, and patient 128 was infected with hvKp), and two patients who were infected with cKp strains that expressed low levels of CPS (appendix p 17).

Using flow cytometry, we measured the convalescent plasma IgG from each patient that bound to their infecting strain or to an isogenic CPS deficient mutant (*ΔwcaJ*) derived from their infecting strain (figure 4). Three of the four study participants demonstrated an increase in IgG to their CPS-deficient mutants compared to the wild-type infecting strain. This shows that a large fraction of the IgG response in patients with Kpn BSI targets non-CPS outer membrane structures, but that these antibodies are blocked from binding by physiologic amounts of CPS produced by hyperencapsulated as well as non-hyperencapsulated strains.

**Figure 4.**
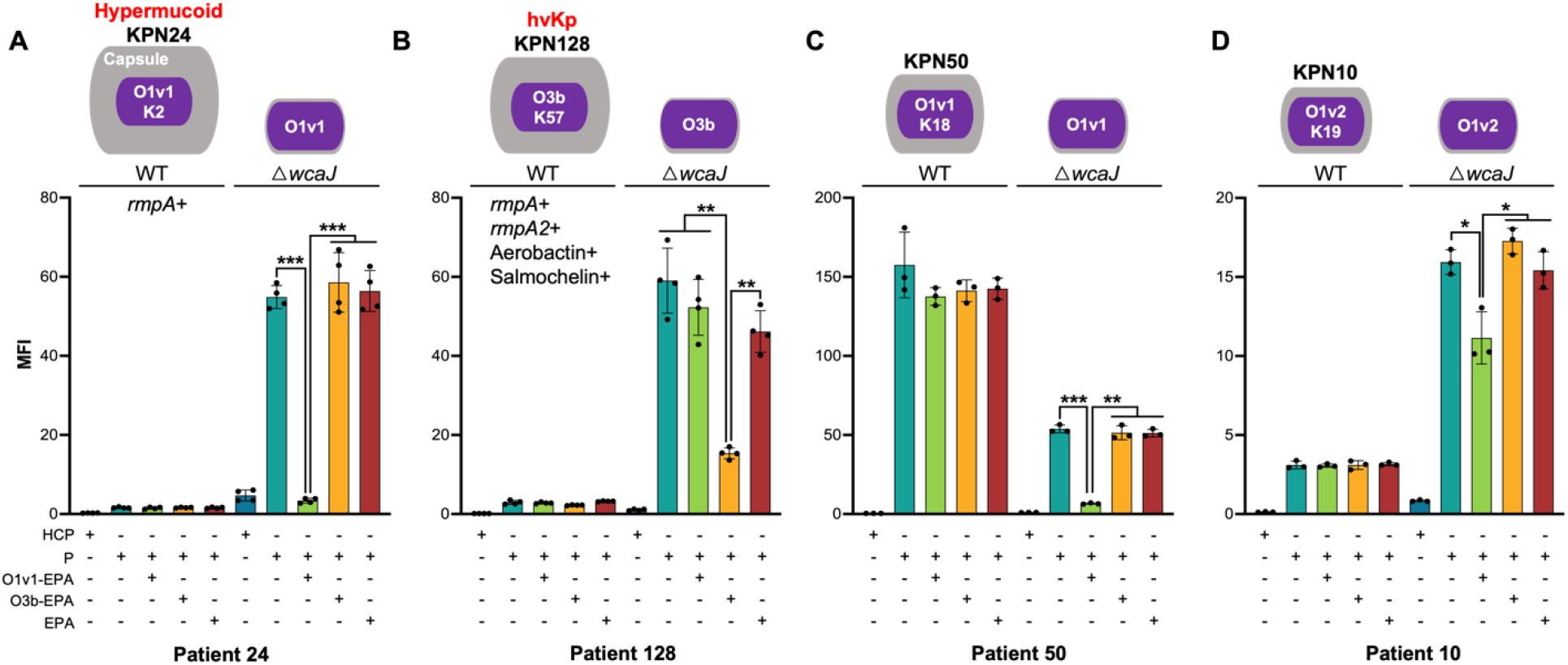
Inhibition of OPS antibody binding by capsule. Pooled plasma sample from 36 healthy individuals (HCP) was used as a baseline reference. Additionally, plasma samples (P) with the highest antibody responses to the OPS expressed by Kpn isolates from patients with Kpn BSI, and those adsorbed with O1v1-EPA, O3b-EPA, and EPA, were analyzed. Specifically, these plasma samples and Kpn isolates were obtained from patients **(A)** 24, **(B)** 128, **(C)** 50, and **(D)** 10. Median fluorescence intensity (MFI) values divided by 1000, representing the quantification of IgG binding to both wild-type Kpn and its capsule-deficient mutant (Δ*wcaJ*), are displayed on the Y-axis. Asterisks indicate statistical significance with P values smaller than 0.05, determined by Welch’s t-test. Significance levels are denoted as follows: * for 0.01 ≤ P < 0.05, ** for 0.001 ≤ P < 0.01, and *** for P < 0.001. The illustration above the figures shows the relative capsule levels in the four Kpn WT strains and their Δ*wcaJ* mutants, based on data presented in appendix (p 17). It also outlines the OPS and CPS serotypes of Kpn isolates as identified by Kleborate analysis in appendix (pp 21-22), with specific characteristics of KPN24 and the hypervirulent (hvKp) KPN128 depicted in the figures. (*N*=4 in (A), (B), *N*=3 in (C), (D))

To determine whether these antibody responses targeted OPS, we then pre-adsorbed the patients’ plasma using OPS. As shown in Figure 4, a significant fraction of the non-CPS targeted IgG antibody response in all 4 of the participants was adsorbed by homologous OPS. Taken together, these data show that OPS is a dominant bacterial cell surface target of the human IgG response to invasive Kpn infection, but that CPS production interferes with OPS-targeted antibody binding.

Similarly, we measured C3 complement deposition following the opsonization of Kpn strains and their corresponding capsule-deficient mutants with purified immunoglobulin (Ig) from each patient. Notably, there was increased antibody-dependent C3 deposition on the capsule-deficient mutants compared to the wild-type infecting strain (appendix p 18). In this experiment, pre-adsorption of Ig with OPS of the serotype matching the target infecting strain, resulted in a significant reduction in C3 deposition on capsule-deficient mutants. These findings confirm the functional capacity of OPS antibodies to facilitate C3 recruitment on Kpn, crucial for opsonization and formation of the membrane attack complex. However, our data underscore a critical interference mechanism, wherein bacterial CPS obstructs this immunological process by concealing OPS antigens from antibody recognition not only in hyperencapsulated strains but also in cKp strains which express lower levels of capsule.

## Discussion

OPS is a candidate vaccine antigen for Kpn^5,14,15^ and is immunogenic in mice and rabbits^24-26^ OPS vaccines have variable efficacy in animal models, with protection dependent on the type of model (e.g. the type of animal, the type of immunization, the route of infection, and the type of challenge strain^9,27-29^). In contrast, there are scant data on the immunogenicity of OPS in humans. To address this knowledge gap, we characterized plasma antibody responses to OPS in a cohort of patients with Kpn BSI. There were three major findings: First, OPS is highly immunogenic in humans with Kpn BSI, independent of capsule expression. Second, there is notable cross-reactivity among similar OPS types in humans. Third, even low levels of capsule expressed by cKp strains interfere with OPS antibody binding and function *in vitro*. These findings have implications for the design of Kpn vaccines.

The primary finding of the study is that OPS is a dominant antigen in human Kpn infection, highlighting its potential as a vaccine component. While this is largely consistent with data from animal models^24,25^, our study did not recapitulate data from mouse models which suggest the O2 serotype may be less immunogenic than other serotypes.^26^ In contrast, we found no evidence that the O2 serotype was less immunogenic in humans with invasive Kpn infection.

Surprisingly, OPS was a dominant target of the antibody response, even in cases of infection with heavily encapsulated Kpn strains. This means that most individuals with Kpn BSI had a response to OPS, and that a large fraction of antibodies which bound to the bacterial cell surface targeted OPS. This contradicted our initial hypothesis that infection with strains expressing high levels of CPS would be associated with reduced antibody responses to OPS. This may be because capsule production does not interfere with OPS antigen presentation in the context of invasive disease (in which bacteria are killed and eliminated by immune cells), or because at certain stages of *in vivo* infection, CPS production may be down-regulated allowing for adequate presentation of OPS antigen on the surface of live bacteria.^30,31^

It is also notable that OPS was immunogenic in a hospitalized population with invasive Kpn. Over half of the patients in our cohort were immunocompromised; nevertheless, OPS antibody responses were relatively preserved in these individuals, compared to responses to the protein antigen MrkA. This may have implications for vaccine development, since most individuals at risk for Kpn have impaired immunity.^4,32^ Whether an OPS-based vaccine, as opposed to natural Kpn infection, could elicit antibody responses in a similar immunocompromised population at risk for invasive Kpn infection would need to be evaluated empirically.

Our study highlighted significant cross-reactivity among related Kpn serotypes, which has crucial implications for vaccine development. The notable cross-reactivity between the O1v1/O1v2 and O2v1/O2v2 serotypes, which mainly differ by the addition of a single galactopyranose in the v2 genotypes, underscores the potential for streamlined vaccine formulations^24,33^. Unlike the limited cross-reactivity observed with the *V. cholerae* O1 Inaba and Ogawa antigens, which differ only by the presence of an additional methyl group on the terminal perosamine in the Inaba serotype and which necessitates the inclusion of both in most cholera vaccines,^34^ the inclusion of both the O1v1 and O1v2 subtypes in a multivalent Kpn vaccine might be unnecessary. The high cross-reactivity observed among key serotypes implies that a Kpn OPS-based vaccine could achieve broad coverage with a select few subtypes, possibly reducing the need for a more extensive serotype representation. This efficiency in serotype selection could simplify vaccine design and improve its protective scope against common Kpn serogroups.

Without established benchmarks of immunity against Kpn infection in humans, we were unable to determine whether individuals in our cohort mounted a protective response against re-infection. However, immunity against Kpn is theorized to involve antibody binding to the bacterial cell, facilitating innate immune system functions such as ADNP or ADCD, which are protective in animal models of Kpn infection.^1,35^ Therefore, it is concerning that OPS antibody-mediated complement deposition on the bacterial cell surface was blocked by CPS, not only in hyperencapsulated strains, but also by lower physiologic amounts of capsule produced by cKp isolates. This finding underscore concerns about the protective capacity of OPS vaccines for invasive Kpn infection.^9^

Our study does have several limitations. First, it included a lower number of patients infected with O2v1, O2v2, O3, and O5 strains relative to those with O1v1, O1v2, and O3b strains, and we did not evaluate antibody responses to less prevalent OPS serotypes, including O4 and O12. Secondly, the investigation does not directly quantify antibody responses to specific CPS serotypes due to the non-dominance of any CPS serotype within our cohort; instead, it examines the influence of CPS by employing CPS-deficient strains. Lastly, the study was restricted to patients with verified BSI. Antibody responses during BSI may not be representative of the full spectrum of end-organ invasive Kpn disease. However, we chose to focus on BSI, since it is a highly specific indicator of invasive infection, while mucosal surface cultures do not distinguish colonization from invasive disease.

In summary, preventing human disease caused by Kpn is critical to the control of AMR bacteria, an urgent global health crisis. Our results demonstrate that OPS is a dominant target of the immune response to invasive Kpn in humans, and that closely related OPS subtypes are cross-reactive. However, capsule might render OPS-targeted antibodies incapable of protecting against infection. These findings have implications for vaccine development targeting this formidable pathogen.

## Contributors

W.H, J.B.H, R.C.L, D.A.R, and P.L.W designed the study. S.E.T, D.S, and K.H.V collected plasma samples and *Klebsiella pneumoniae* isolates. E.O, D.J.R, R.C.L, and J.B.H analyzed clinical data of patients and healthy individuals. D.S, S.S, and K.H.V conducted whole genome sequencing and analysis. W.H, D.S, K.H.V, B.B, J.Z, and P.L.W performed the experiments. W.H and S.R.R analyzed and visualized the data. A.S.C, C.M.H, M.A, and C.J.K constructed and provided HSA- and EPA-conjugated OPS. W.H and J.B.H wrote the original draft. All authors reviewed and edited the manuscript.

## Data sharing

All data and materials necessary to reproduce the findings of this study are comprehensively documented within the main paper and its supplementary materials. The plasmids constructed and utilized in this research are detailed in the supplementary materials, and the primer sequences employed can be found in appendix (pp 24-25).

## Acknowledgements

This study was supported by National Institute of Allergy and Infectious Diseases grant R01AI175345 (to J.B.H.).

