## Supplementary data for "Antibody responses in *Klebsiella pneumoniae* bloodstream infection: a cohort study"

Contents:

Supplementary Methods (pages 3 to 9)

Figure S1 Flowchart illustrating the patient selection process for individuals with *Klebsiella pneumoniae* bloodstream infection (page 10)

Figure S2 Longitudinal antibody responses to less dominant OPS in patients with Kpn BSI (page 11)

Figure S3 Correlation between capsule amount of infecting strain and OPS antibody response (page 12)

Figure S4 Preserved OPS antibody response in immunocompromised patient (pages 13 to 14)

Figure S5 Antibody responses to OPS and MrkA protein in patients with Kpn BSI (page 15)

Figure S6 OPS antibody function (page 16)

Figure S7 Capsule quantification of Kpn isolates and capsule knockout mutants (page 17)

Figure S8 OPS antibody mediated complement deposition on Kpn isolates (page 18)

Figure S9 MrkA homologues in 69 Kpn isolates (page 19)

Figure S10 Adsorbed OPS antibody by adding OPS and heat-killed Kpn isolates (page 20)

Table S1 Kleborate results of 69 *Klebsiella pneumoniae* isolates (pages 21 to 22)

Table S2 The number of plasma samples used in Figure 2, Figure 3B, and Figure S5 (page 23)

Table S3 Primers used in this study (pages 24 to 25)

Supplementary References (page 26)

### Material and Methods

**Enrollment of patients with *Klebsiella pneumoniae* blood stream infection and controls**

This investigation was carried out at Massachusetts General Hospital (MGH) a 1000-bed tertiary care hospital which provides care for patients of all ages, with approval from the MassGeneral Brigham Institutional Review Board. We enrolled a cohort of all sequentially identified patients with *Klebsiella pneumoniae* (Kpn) blood stream infection (BSI), as identified by the clinical microbiology laboratory according to Clinical and Laboratory Standards Institute Guidelines, over a one-year period from 07/24/21 to 08/04/22. Demographic and clinical data were extracted from the medical record, as described previously.[1] Excess plasma used for this study was collected longitudinally for routine patient care over the course of each patients' hospitalization or on subsequent follow-up visits. We excluded patients who had inadequate plasma collected due to low sample volume, death or lack of follow-up visits (Figure S1). We excluded patients whose isolates were not confirmed to be *Klebsiella pneumoniae* by subsequent whole genome sequencing. We compared immune responses in the KPN BSI cohort with a previously described cohort healthy adults (HC) presenting for a routine outpatient pre-travel consultation at MGH[2], and with a cohort of patients with *Enterococcus* spp. BSI (BC) hospitalized at MGH. One plasma sample per each healthy adult was collected. For patients with *Enterococcus* spp. BSI, a single plasma specimen between day 7 and day 14 following their first positive blood culture was collected. An immunocompromised patient was defined as having a history of acquired immunodeficiency syndrome, a rheumatologic condition requiring specific immunosuppressive treatment within the past 3 months, cancer actively undergoing chemotherapy within the past 3 months, being a solid organ transplant recipient with ongoing immunosuppressive therapy, or having primary immunodeficiency.

**Whole genome sequencing (WGS) and analysis**

We performed WGS on Kpn isolates as described previously.[1] In brief, gDNA was sequenced at the Vanderbilt University Medical Center Vantage core on an Illumina NovaSeq6000 system. Sequenced isolates yielded an average of 10 million reads. Adapter-trimmed sequences were assembled using SPAdes[3] and Kleborate was used to confirm Kpn identification and for *in silico* OPS and CPS serotyping.[4] Hypervirulent Kpn (hvKp) was classified based on the presence of either a complete *rmpA* or *rmpA2* gene in conjunction with complete aerobactin and salmochelin genes (Table S1).[5] BlastN analysis involved the comparison of *wbbY* reference sequence from the Kleborate database with the whole genome assemblies of O2v1 and O2v2 KPN isolates. If fragmented *wbbY* genes were distributed across different contigs in these isolates, the OPS serotype was reclassified as O1v1 or O1v2. The

Prokka tool[6] was utilized to identify open reading frames (ORFs) in the 69 whole genome assemblies of Kpn isolates. BlastP analysis of ORFs and the MrkA reference sequence was performed, with >90% identity and >85% gene length coverage result considered a homologous protein of MrkA. MrkA homologous proteins were clustered using the CLUSTALW web-based tool [7] and visualized with Jalview[8] (Figure S9). In the comparison of MrkA homologues, KPN6 and KPN55, lacking a MrkA homologue protein, were excluded from the analyses of the MrkA antibody response. For constructing *wcaJ* gene knockout mutants, WcaJ reference sequences of each CPS type (KPN10-K19, KPN24-K2, KPN50-K18, KPN128-K57) from the Kleborate database were used. BlastP searches with WcaJ reference sequences and ORFs of the four Kpn isolates were conducted, and the protein with >98% identity and >98% gene length coverage was identified as the WcaJ homologue. The Artemis tool[9] was employed to extract nucleotide sequences upstream and downstream of the WcaJ homologue for constructing a Δ*wcaJ* mutant.

**Antigen preparation**

O-antigens conjugated to Exoprotein A of *Pseudomonas aeruginosa* (EPA), as well as EPA itself, were prepared following established protocols.[10] O-specific polysaccharides (OPS) conjugated to Human Serum Albumin (HSA) were synthesized as follows: A mixture containing OPS, HSA, and sodium cyanoborohydride in a ratio of 3:1:6 (w/w/w) was stirred in 0.1 M borate buffer at pH 7.4 and maintained at 55°C for seven days, followed by dialysis in water. The MrkA protein (LSBio) served as the protein antigen for Kpn. EPA and HSA (Sigma) antigens were employed as controls for the bioconjugated antigens.

**Multiplexed bead assay (MBA)**

Antigens were conjugated to magnetic beads according to the xMAP® Antibody Coupling Kit (Diasorin) recommended protocol. Plasma from HC, BC and patients with Kpn BSI were diluted 1:100 in phosphate buffed saline (PBS) containing 1% BSA with 0.05% Tween20, and 5μl of the diluted plasma was added to 384-Well Polystyrene Non-Binding Flat Bottom Microplate (Fisher Scientific Greiner Bio-One™) and incubated with the antigen coated bead mixture at room temperature for 2 hours, then washed three times. Subsequently, PE-conjugated mouse anti-human IgG, IgM, and goat anti-human IgA secondary antibodies (Southern Biotech) were added to each well and incubated for 1 hour. Following the incubation, the coated bead mixture was washed and reconstituted in sheath fluid (Fisher Scientific). Fluorescence data from each well was acquired on a Luminex FlexMap3D. A standard dilution series, prepared from a mixture of patient serum, was run independently on 4 experimental plates. Median Fluorescence Intensity (MFI) values were normalized across 4 individual MBAs by comparing standard curves. Specifically, the coefficient of variation (CV) of each antigen at each dilution factor between 4 standard curves was calculated. The 4 MFIs at the dilution factor with the lowest CV were used to normalize MFIs inter MBAs. A small number of plasma samples exhibited robust antibody responses to the EPA antigen. Plasma samples with MFI to the EPA antigen exceeding the 75th percentile (Q3) plus 1.5 times the interquartile range (IQR) of 36 healthy controls (calculated as Q3 + 1.5IQR) were identified as EPA-positive and subsequently excluded from the results involving EPA-conjugated bead. In Figure 3A, plasma samples with MFIs lower than 200 for HSA and EPA antigens were included. This approach was implemented to mitigate potential bias arising from varying antibodies binding to different conjugative proteins. 29 (7 HC, 6 BC, and 16 patients with Kpn BSI), 30 (3 HC, 3 BC, and 24 patients with Kpn BSI), and 35 (9 HC, 4 BC, and 22 patients with Kpn BSI) human plasmas were retained for IgG, IgM, and IgA analyses, respectively. All results from the MBAs were visualized using seaborn (v0.11.2) and matplotlib (v3.4.3) in Python (v3.9.7).

**Antibody-dependent neutrophil phagocytosis (ADNP)**

Plasma collected the closest to day 10 after the first positive Kpn blood culture from patients with O1v1, O1v2, and O3b Kpn BSI was used to measure ADNP responses. Additionally, 36 plasma samples from healthy controls were included as a comparator group. OPS was modified with DMTMM and coupled to carboxylated fluorescent beads (Thermo Fisher). To form immune complexes, each separately antigen-coupled bead was incubated for 2 hours at 37°C with diluted samples (1:200) and then washed to remove unbound immunoglobulins. The immune complexes were incubated for 1 hour with fresh blood neutrophils isolated from healthy donors with a commercially available kit (StemCell). Following the incubation, cells were fixed with 4% paraformaldehyde and flow cytometry was performed to identify the percentage of cells that had phagocytosed beads as well as the number of beads that had been phagocytosed (phagocytosis score = % positive cells × Median Fluorescent Intensity of positive cells/10000). Flow cytometry was performed with an IQue (Intellicyt), and analysis was performed on IntelliCyt ForeCyt (v8.1) or using FlowJo V10.7.1.

**Antibody dependent complement deposition (ADCD)**

The same plasma samples used in the ADNP assay were utilized for ADCD.  ADCD was conducted using a 384-well based customized multiplexed assay. OPS antigens were modified by 4-(4,6-dimethoxy[1,3,5]triazin-2-yl)-4-methyl-morpholinium and conjugated to Luminex Magplex carboxylated beads. To form immune complexes, a mixture of antigen-coupled beads was incubated for 2 hours at 37°C with diluted samples (1:200) and then washed to remove unbound immunoglobulins. Lyophilized guinea pig complement (Cedarlane) was resuspended according to manufacturer's instructions and diluted in gelatin veronal buffer with calcium and magnesium (Boston BioProducts). Resuspended guinea pig complement was added to immune complexes and incubated for 20 minutes at 37°C. Post incubation, C3 was detected with Fluorescein-Conjugated Goat IgG Fraction to Guinea Pig Complement C3 (Mpbio).

**Bacterial strains and growth conditions**

All Kpn isolates and *Escherichia coli* (*E. coli*) strains utilized in cloning experiments were cultured in Luria-Bertani (LB) medium with agitation at 200 rpm. Antibiotic-selective media, namely LB supplemented with apramycin (Apr), spectinomycin (Spec), and carbenicillin (Carbe), were employed for genetic manipulation and bacterial growth carrying antibiotic-resistant plasmids. The antibiotic susceptibility profile for the four Kpn isolates (KPN10, KPN24, KPN50, and KPN128) included susceptibility to 30μg/ml Apr and 50~100μg/ml Spec, and *E. coli* S17-1 λpir was susceptible to 50μg/ml carbenicillin. Single bacterial colonies were selected from LB agar plates or LB agar supplemented with antibiotics and inoculated into LB or LB with antibiotics for precultures, grown overnight at 30℃ or 37℃ with shaking at 200rpm. Precultures were then diluted 100-fold for subcultures and incubated similarly to the preculture conditions.

**Plasmid construction (pKPGFP, pDMS197-apr)**

The plasmids pKPGFP and pDMS197-apr were engineered for this study. The backbone of pKPGFP was derived from pBad24-sfGFPx1 (addgene, #51558). In the pKPGFP plasmid, the *araC* gene was removed, and the *araBAD* promoter sequence was replaced with the *rpsL* promoter from pCasKP-apr (addgene, #117231) to facilitate endogenous GFP expression. The ampicillin-resistant gene (*ampR*) in pKPGFP was substituted with the apramycin-resistant gene from pCasKP-apr. pDMS197-apr utilized the pDMS197 plasmid backbone (addgene, #43831). In this plasmid, the tetracycline promoter and its corresponding resistance gene were replaced with the *ampR* promoter and apramycin-resistant gene from pKPGFP. Both plasmids were constructed using the Gibson assembly process. The primer design for the fragments for the two plasmids was facilitated through the NEBuilder Assembly Tool. These fragments were amplified with Q5 High-Fidelity DNA polymerase (NEB), and the linear PCR products were ligated using NEBuilder HiFi DNA assembly master mix (NEB). Subsequently, pKPGFP was transformed into DH5alpha chemically competent cells (Invitrogen), and pDMS197-apr was transformed into One Shot PIR1 chemically competent *E. coli* (Invitrogen). The transformed *E. coli* containing these plasmids were selected on LB agar with Apr.

**_wcaJ_ deletion**

To generate capsule-deficient mutants of three Kpn isolates with O1 serotypes (KPN10, KPN24, KPN50), we used the CRISPR-Cas9 and Red recombineering systems as described elsewhere.[11] Briefly, the *wcaJ* homologue gene of the Kpn isolates was input into the CRISPRdirect website-based tool[12] to identify a suitable single guide RNA (sgRNA) sequence. The sequence-confirmed pSGKP-spec harboring sgRNA and homologue sequences for repairing sequence were co-transformed into electrocompetent Kpn isolates harboring pCasKP-apr. The repairing sequences included a synthetic 90-mer double-stranded DNA composed of each 45-mer of both upstream and downstream sequences of the *wcaJ* gene in constructing KPN50Δ*wcaJ* and approximately 1.3 kb homologue sequences composed of both ~0.65 kb upstream and ~0.65 kb downstream sequences of the *wcaJ* gene in constructing KPN10Δ*wcaJ* and KPN24Δ*wcaJ*. After incubating transformants on LB agar with Apr and Spec at 30℃, Sanger sequencing was performed to confirm the sequence mismatch and identify Δ*wcaJ* mutants, following which pCasKP-apr and pSGKP-spec harboring sgRNA were cured.

Because this method failed to generate a *wcaJ* knockout mutant of KPN128, we used an alternative approach involving a two-step allelic exchange method using a modified pDMS197-apr plasmid. pDMS197-apr harboring flanking regions (~0.65 kb homologue sequences upstream and downstream of the *wcaJ* gene), pDMS197-apr_KPN128*wcaJ*, was electroporated into electrocompetent *E. coli* S17-1 λpir. Exponential-phase cultures of KPN128 and S17-1 λpir containing pDMS197-apr_KPN128*wcaJ* were spread on LB agar and LB agar with Apr overnight. Subsequently, bacterial mating was incubated for 5 hours on LB agar, followed by plating on LB agar with Apr and Carbe to identify chromosomally integrated pDMS197-apr_KPN128*wcaJ* in KPN128 (single-crossover events). Merodiploid colonies were streaked on no-salt LB supplemented with 15% sucrose at room temperature for over 24 hours to induce *sacB*-mediated plasmid removal (double-crossover events). The identification of the deletion mutant was achieved through colony PCR, and Sanger sequencing was performed to confirm the gene deletion in the KPN128.

**Bacterial flow cytometry**

Four Kpn isolates (KPN10, KPN24, KPN50, KPN128) harboring the pKPGFP plasmid were obtained through electroporation and streaked on LB agar + Apr. The overnight-grown cells were washed with PBS containing 20%

glycerol, and aliquots of cells with a final OD of 1 were stored at -80°C for use in subsequent experiments. Plasmas exhibiting the highest antibody responses to the O1 antigens from patients with O1 Kpn BSI (patients 10, 24, and 50) and to the O3b antigen from patient with O3b Kpn BSI (Patient 128) were selected for this experiment. A pooled plasma sample consisting of 36 plasmas from healthy individuals served as a control plasma. Plasma complement was inactivated at 56°C for 30 minutes. For pre-adsorbed plasma, diluted plasma in flow buffer (2% BSA in PBS) was incubated with O1v1-EPA, O3b-EPA, and EPA antigens. Subsequently, 25µL of GFP-labeled bacteria and 25µL of diluted plasma or pre-adsorbed plasma were added to a V-bottom 96-well plate. The final bacterial CFU in this reaction was approximately 1.5E+6, the final dilution factors of plasma were 1:400 for patient 24's plasma and 1:1600 for patients 10, 50, and 128's plasma. After incubating at 37°C for 15 minutes with shaking at 200rpm, 200µL of flow buffer was added, and bacteria were spun at 4,000g for 10 minutes at 4°C and washed. Subsequently, 50µL of 1:100 diluted goat anti-human IgG-PE secondary antibody (Southern Biotech) was added and resuspended by pipetting. After an incubation at room temperature, a second washing step with flow buffer was performed. The samples were then resuspended in flow buffer and the plate was read on a BD CSampler Plus to collect at least 20,000 events. The threshold of FSC-H was set at 9,000, and fluidics speed was controlled not to exceed 2000/sec. Data was analyzed using FlowJo v10.9.0, and median MFIs of PE fluorescence were extracted after gating GFP-positive Kpn.

C3 deposition on Kpn isolates was evaluated utilizing Gelatin Veronal Buffer (GVB; Sigma). Immunoglobulins (Ig) from the previously mentioned plasma samples underwent were purified using NAb Protein A/G Spin Columns (Thermo Scientific). These Ig, at concentrations of 20µg/ml for samples from patients 10 and 128 and 10µg/ml for samples from patients 24 and 50, along with Ig pre-adsorbed with O1v1-EPA, O3b-EPA, and EPA antigens, were used for bacteria opsonization. Following this, 50µL of diluted guinea pig complement (Sigma) was incubated with the opsonized GFP-labeled bacteria at 37°C for 30 minutes with agitation at 200rpm. A subsequent washing step was performed using flow buffer before the addition of 50µL of 1:3200 dilution of goat anti-complement C3 polyclonal IgG (Invitrogen) and incubation at room temperature on a microplate shaker. After another washing step, 50µL of a 1:1000 dilution of rabbit anti-goat IgG conjugated with Alexa Fluor® 647 (abcam) was added to the samples, which were then incubated again. Following a final wash in flow buffer, flow cytometric analysis was conducted under the same conditions as the previous bacterial flow cytometry experiment. Median MFIs of APC fluorescence were extracted after GFP-positive Kpn were gate.

**Blocking OPS antibody**

To block OPS antibodies, diluted plasma samples in flow buffer and purified Ig in GVB were incubated with antigens (O1v1-EPA, O3b-EPA, and EPA, final antigen concentration: 10μg/ml) at 37°C for 30 minutes with shaking at 200rpm. These pre-adsorbed plasmas and purified Ig were then used in bacterial flow cytometry. As an indirect means to assess OPS production levels in each Kpn isolate and Δ*wcaJ* mutants, Kpn isolates harboring the pKPGFP plasmid were heat-killed (HK) at 100°C for 15 minutes. Subsequently, 25 μL of HK bacteria (1.5E+6 CFU) and 25 μL of diluted plasma were incubated at 37°C for 30 minutes under 200rpm. The final dilution factors of plasmas were 1:400 for patient 24 and 1:1600 for patients 10, 50, and 128. Pre-adsorbed plasmas by antigens or HK bacteria were used for multiplexed bead assays in the same manner as described in the multiplexed bead assay method to assess OPS antibodies were adsorbed (Figure S10).

**Capsule quantification**

Glucuronic acid quantification was performed as described previously.[13] In brief, Kpn isolates and their Δ*wcaJ* mutants, cultured overnight, were concentrated and normalized to an OD600nm ranging from 3.00 to 3.50. After treatment with Zwittergent, the cultures underwent incubation at 50°C for 20 minutes, followed by centrifugation. The supernatant was mixed with cold ethanol, leading to capsule pellet formation upon centrifugation. After resuspension in MilliQ water, a glucuronic acid standard dilution series was prepared. Sulfuric acid was added to each sample and standard, followed by incubation and cooling. Subsequently, 0.15% 3-hydroxydiphenol in 0.5% NaOH was introduced, and the solutions were spectrophotometrically measured at OD520nm. Capsule concentrations were determined by comparing sample OD measurements to the glucuronic acid standard curve.

**Figure S1. Flowchart illustrating the patient selection process for individuals with *Klebsiella pneumoniae* bloodstream infection**

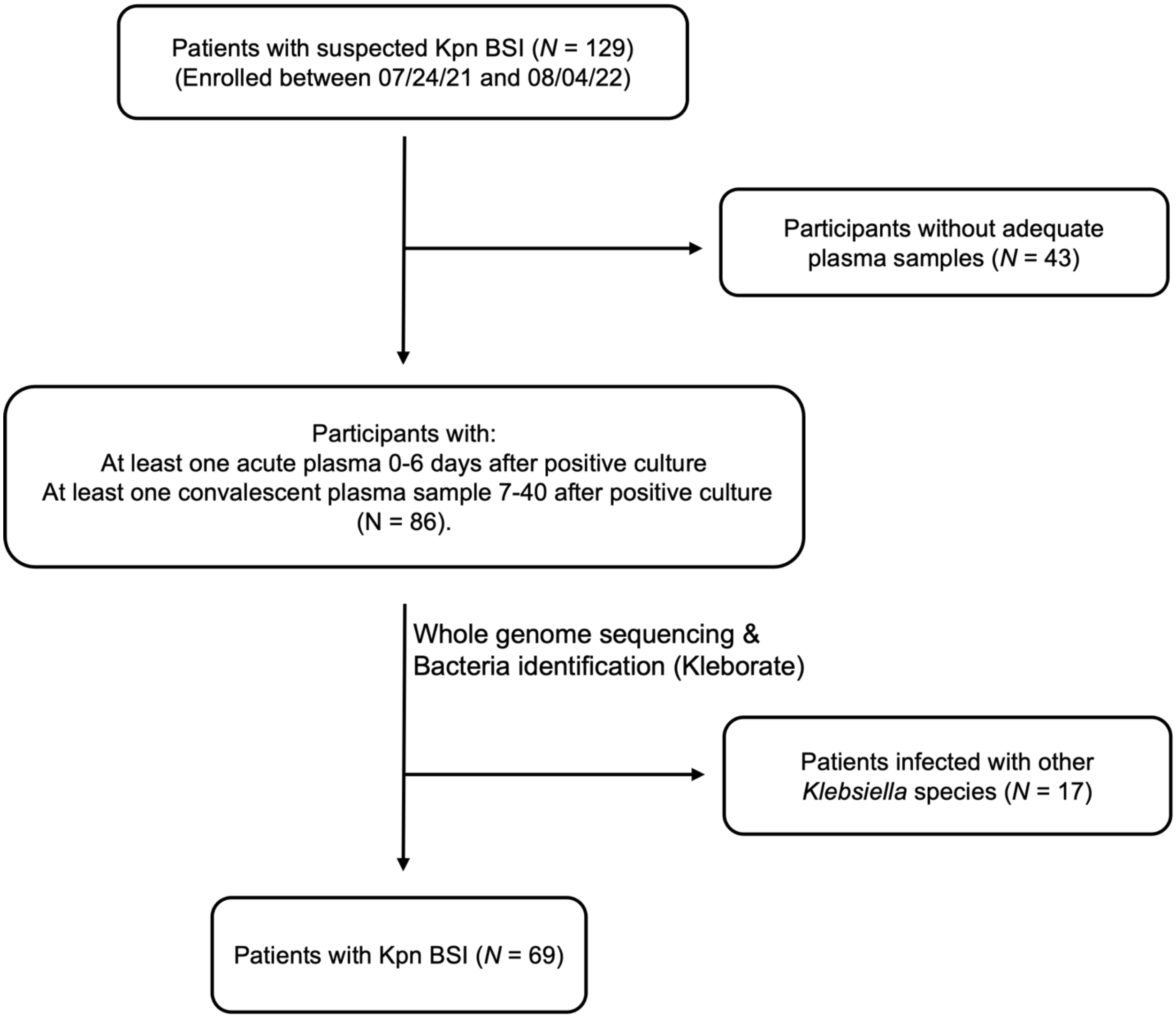

**Figure S2. Longitudinal antibody responses to less dominant OPS in patients with Kpn BSI**

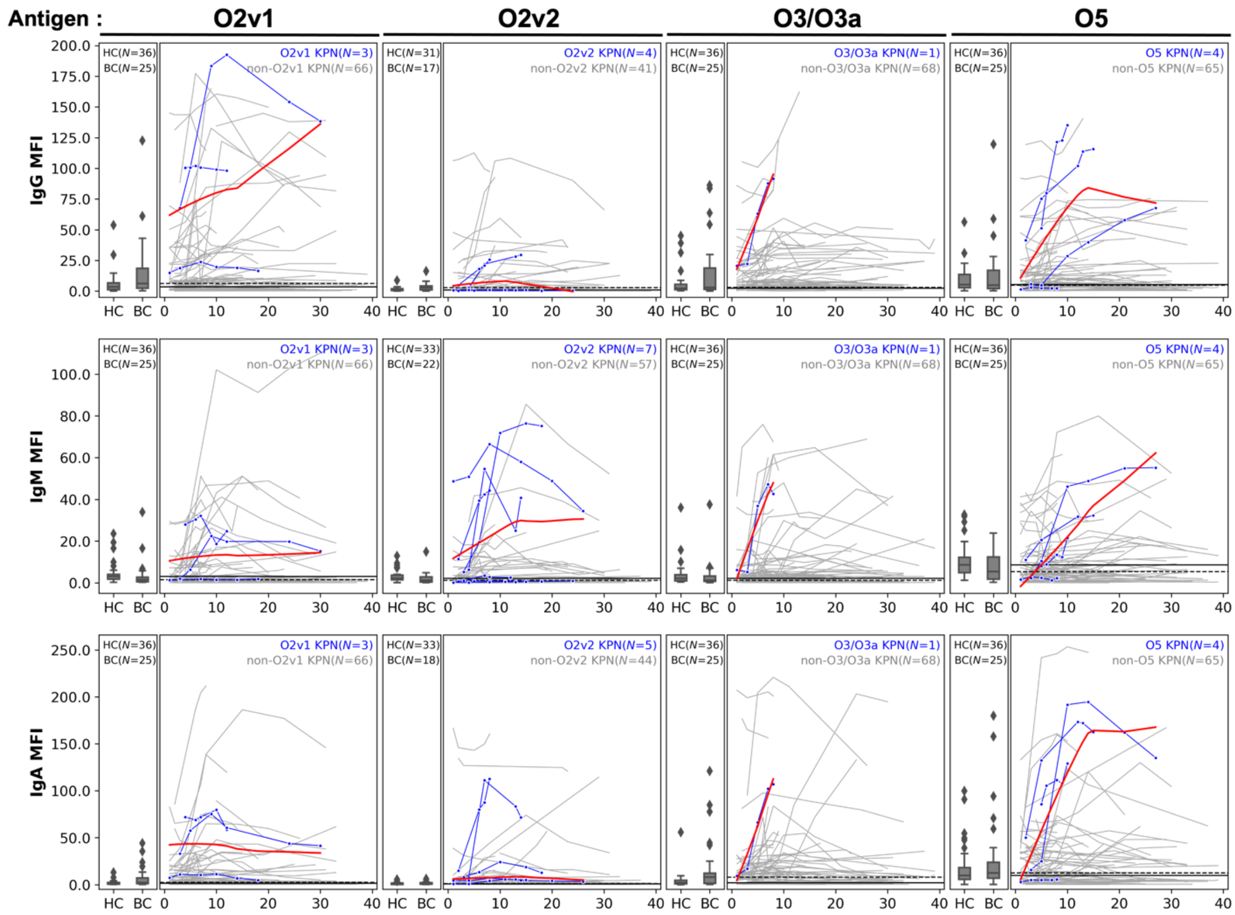


IgG, IgM, and IgA antibody responses to O2v1, O2v2, O3/O3a (O3), and O5 antigens were presented. The Y-axis represents Median Fluorescence Intensity (MFI), divided by 1000. Boxplots compare MFIs to antigens in plasma samples from healthy- (HC) and *Enterococcus* spp. bacteremic-controls (BC). Longitudinal antibody responses in patients with Kpn BSI are represented by individual lines next to boxplots, with each data point corresponding to samples from a single patient at different time points. X-axis denotes the date of plasma collection from patients with Kpn BSI after the first positive blood culture of Kpn (Day 0). Blue lines indicate antibody responses of patients bacteremic with Kpn isolates having the homologous OPS compared to the antigen, while gray lines represent patients bacteremic with Kpn isolates with heterologous OPS. Red lines illustrate the LOWESS regression applied to the blue lines. The solid black line represents the median of HC, and the dashed black line represents that of BC. The number of patients included in each analysis is shown in each graph.

**Figure S3. Correlation between capsule amount of infecting strain and OPS antibody response**

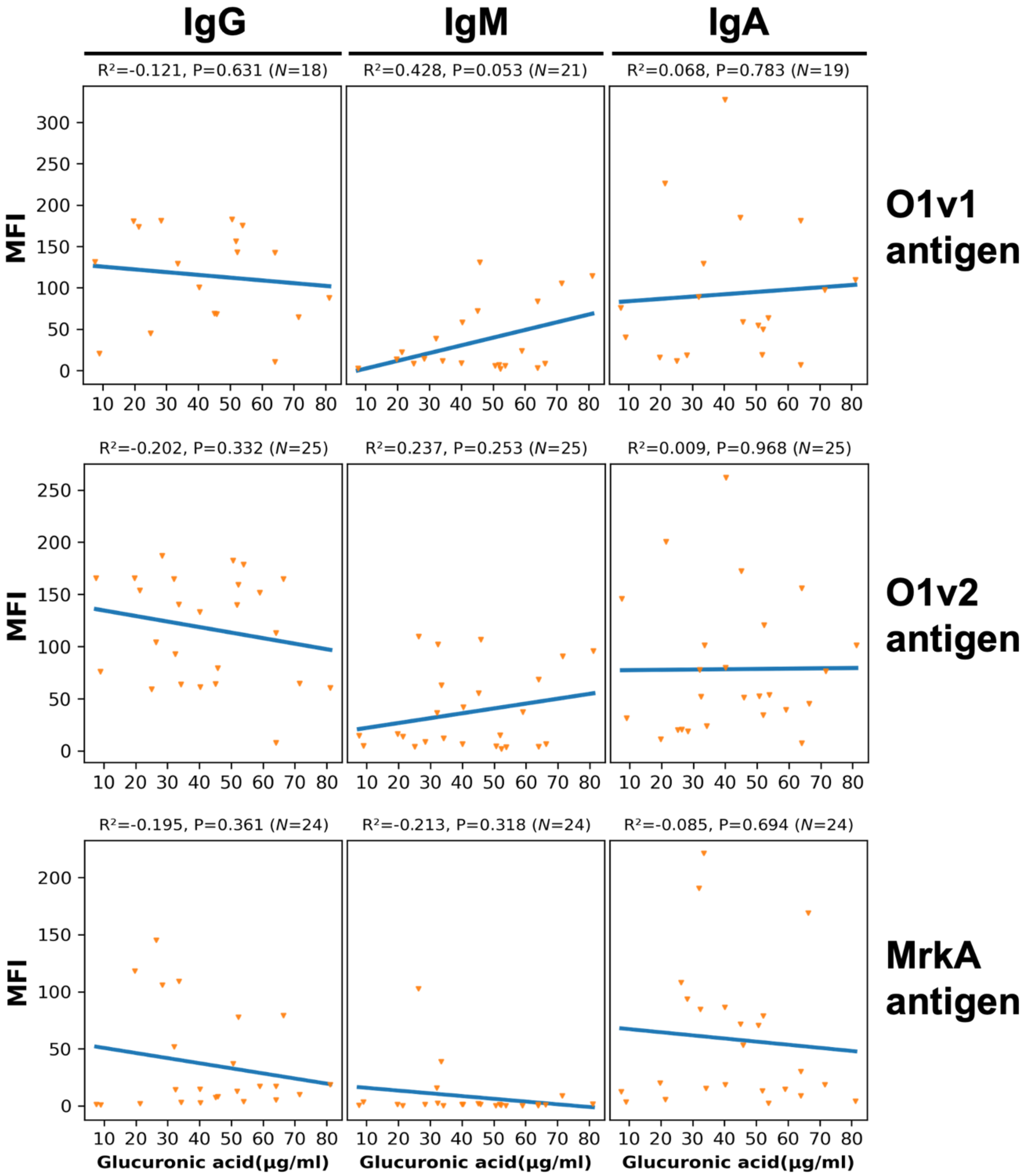


This figure illustrates the correlation between IgG, IgM, and IgA antibody responses (MFI divided by 1000) to O1v1, O1v2, and MrkA antigens in patients with O1 Kpn BSI, and the glucuronic acid levels (indicative of capsule amount) in the associated O1 Kpn isolates. The figure provides the R-squared ($R^2$) values, P values (P) obtained from Pearson correlation analysis, and the number of samples used in the analysis.

**Figure S4. Preserved OPS antibody response in immunocompromised patient**

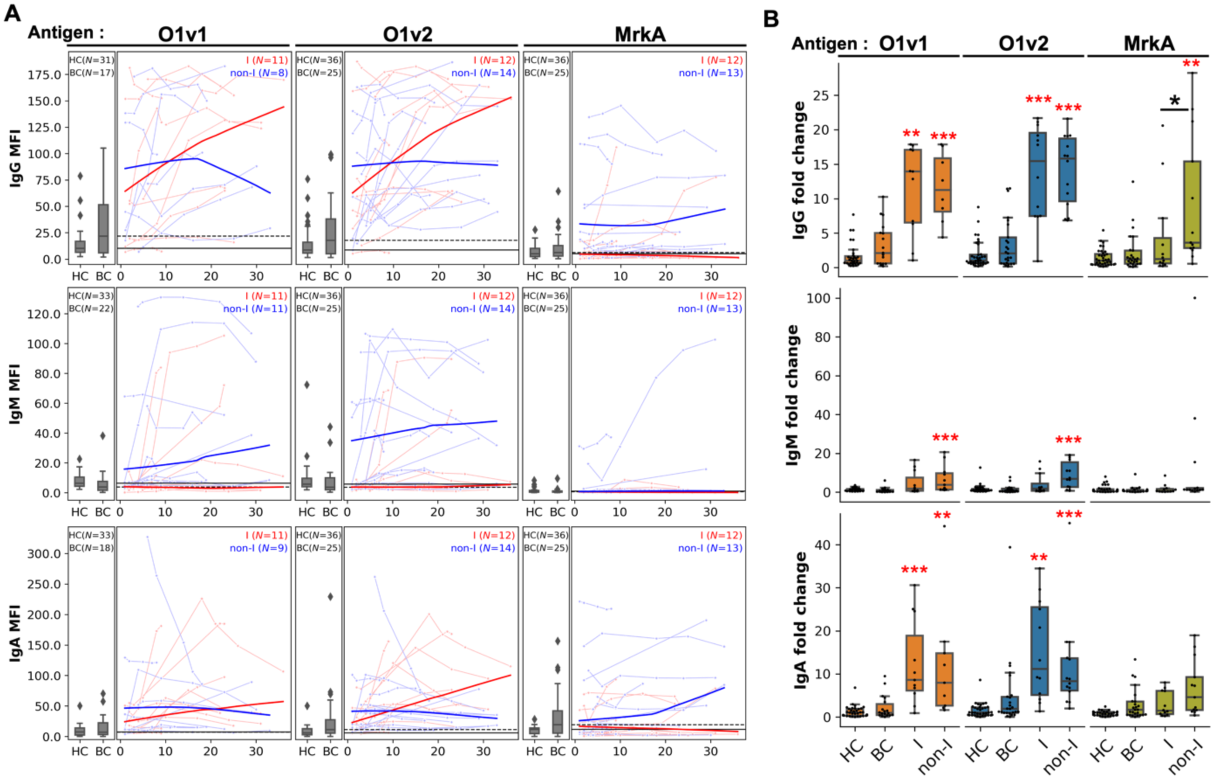


**(A)** O1v1, O1v2 and MrkA responses in plasma samples from healthy controls (HC), patients with *Enterococcus* BSI (BC), and patients with O1 Kpn BSI, were measured. The Y-axis presents MFI values, and each boxplot displays antibody responses to antigens from HC and BC, while longitudinal antibody responses in patients with O1 Kpn BSI are depicted by individual lines next to boxplots. Each data point on a line corresponds to samples from a single patient at various time points. The X-axis represents the date when plasma was collected post the first positive blood culture for Kpn (Day 0). Results from patients with O1 Kpn BSI were categorized into immunocompromised patients (I) and non-immunocompromised patients (non-I). Red lines indicate antibody responses from I, blue lines represent antibody responses in non-I. Darker lines show the result of LOWESS regression applied separately to the I and non-I individuals, highlighting the trend in antibody responses over time. The solid black line represents the median MFI of HC, and the dashed black line represents that of BC. The number of patients for each analysis is shown in each graph. **(B)** The peak antibody responses to antigens from patients with O1 Kpn BSI are shown in fold change units relative to the median MFI of HC. Results are categorized into antibody responses from I and non-I groups. Red asterisks denote P values smaller than 0.05 when compared to both HC and BC, and black asterisks indicate comparisons of fold change in I and non-I groups by Mann-Whitney U test: * $0.01 \le P < 0.05$, ** $0.001 \le P < 0.01$, *** $P < 0.001$. The number of samples is the same as those in (A).

**Figure S5. Antibody responses to OPS and MrkA protein in patients with Kpn BSI**

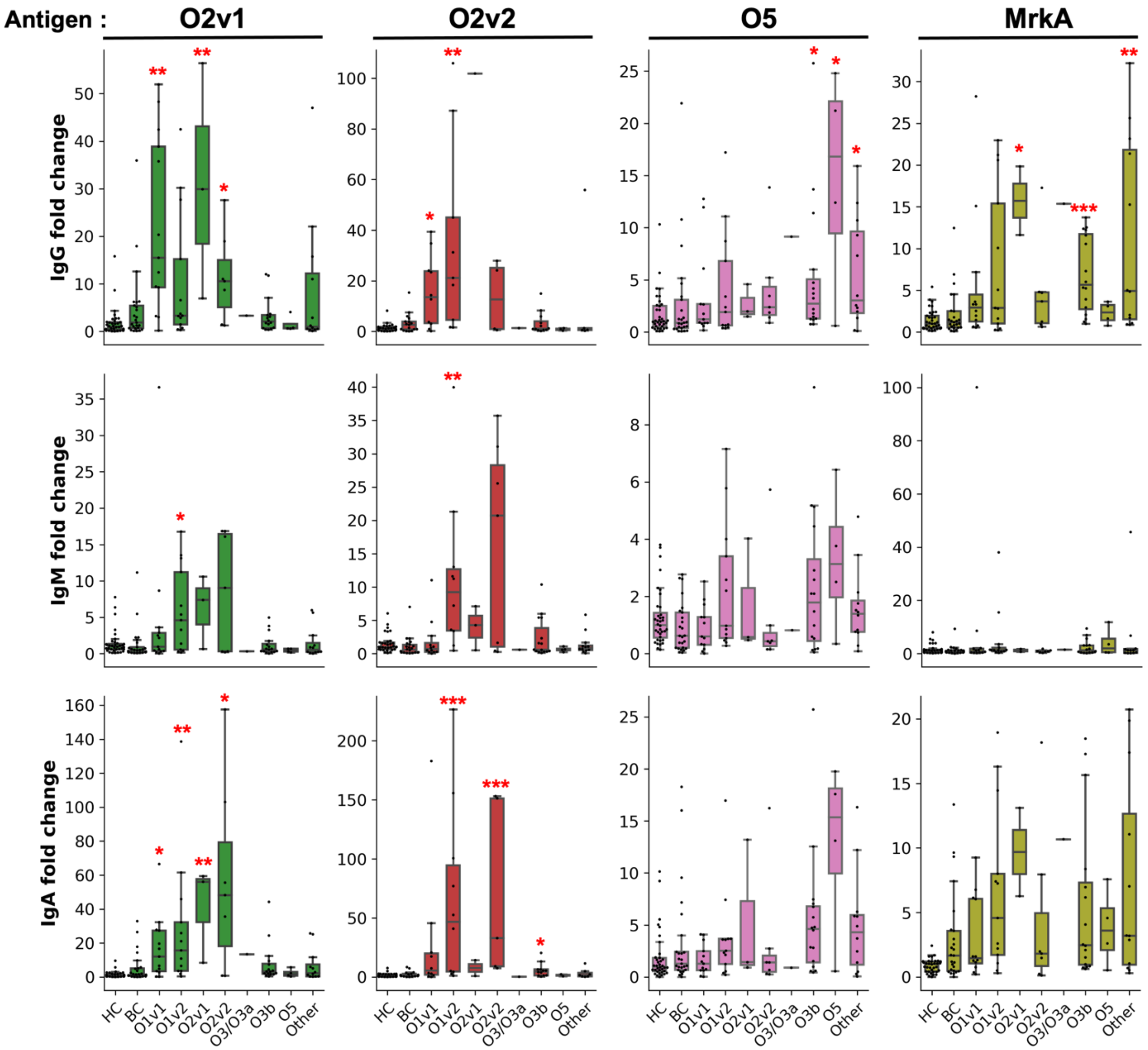


Individual boxplots provide IgG, IgM, and IgA antibody responses for healthy individuals (HC), bacteremic patients with *Enterococcus* spp. BSI (BC), and the highest antibody responses observed in patients with Kpn BSI in response to O2v1, O2v2, O5, and MrkA antigens. The X-axis illustrates plasma samples sourced from HC, BC, and patients infected with Kpn with diverse OPS types. The "Other" category encompasses OPS types O4, O12, and unidentified OPS. The Y-axis represents the fold change in antibody responses relative to HC, calculated as the MFI divided by the median MFI of HC. Red asterisks signify statistical significance (P values < 0.05) when compared to both HC and BC using Mann-Whitney U test. Significance levels are denoted as follows: * for 0.01 ≤ P < 0.05, ** for 0.001 ≤ P < 0.01, and *** for P < 0.001. Detailed information regarding the number of samples is available in Table S3.

**Figure S6. OPS antibody function**

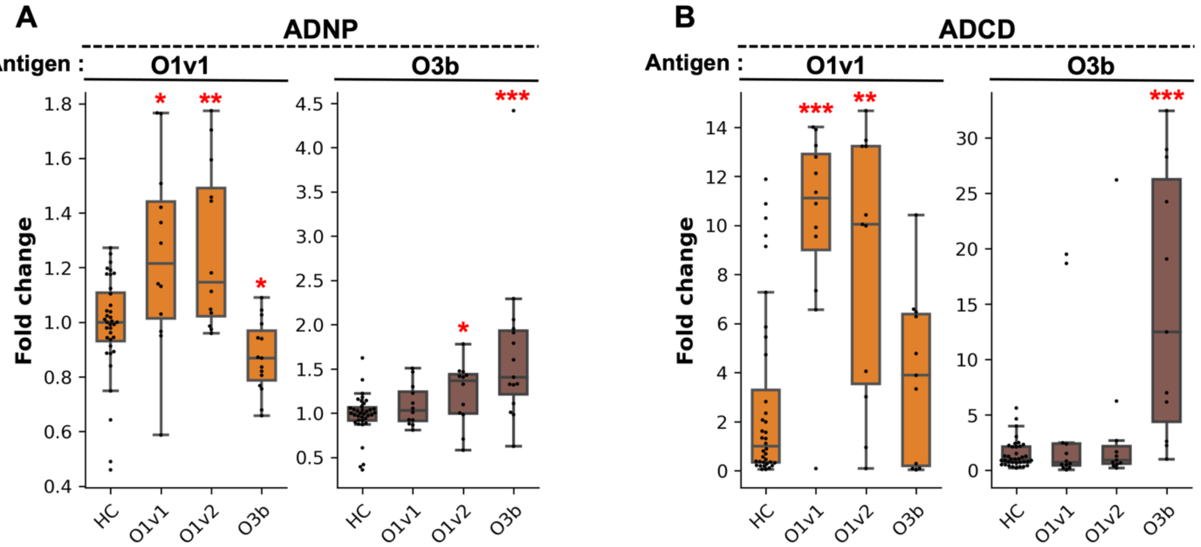


**(A)** ADNP (Antibody Dependent Neutrophil Phagocytosis) was tested using plasma samples from healthy individuals (HC) and patients with Kpn BSI featuring O1v1, O1v2, and O3b serotypes. The ADNP activity of O1v1 and O3b-conjugated beads was normalized by dividing it by the median ADNP activity in HC. The average fold change of ADNP activities, derived from two healthy neutrophil donors, is presented (N=36 HC, 12 O1v1, 12 O1v2, 15 O3b) **(B)** ADCD (Antibody Dependent Complement Deposition) assays were performed using opsonized beads conjugated with O1v1 and O3b serotypes, along with plasma from HC and patients with Kpn BSI presenting O1v1, O1v2, and O3b serotypes. Additionally, guinea pig complement was used in the assays. ADCD activities were divided by the median ADCD activity in HC, and the results are expressed as fold change (N=36 HC, 12 O1v1, 11 O1v2, 11 O3b). Asterisks in both (A) and (B) indicate P values smaller than 0.05 compared to HC by Mann-Whitney U test, with significance levels denoted as follows: * $0.01 \leq P < 0.05$, ** $0.001 \leq P < 0.01$, *** $P < 0.001$

**Figure S7. Capsule quantification of Kpn isolates and capsule knockout mutants**

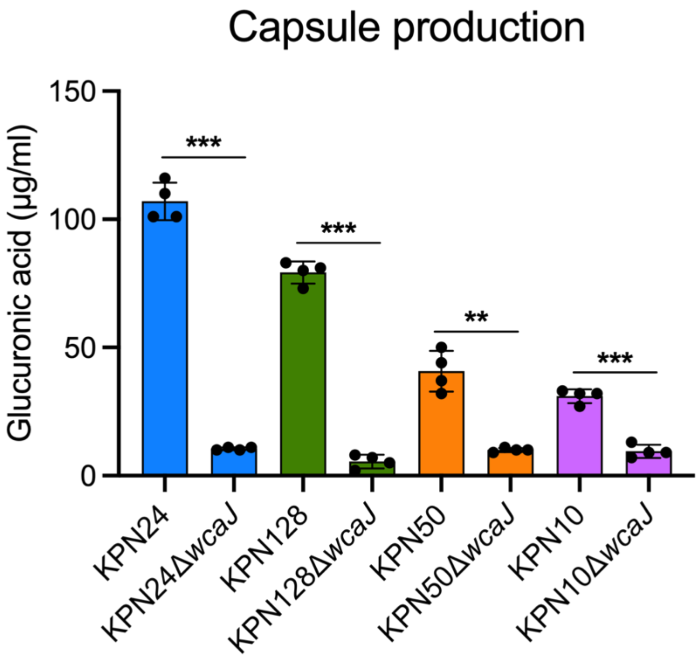


Glucuronic acid levels were quantified using four Kpn wild-type isolates (KPN24, KPN128, KPN50 and KPN10) from patients 24, 128, 50 and 10 and their capsule knockout mutants (Δ*wcaJ*) (*N*=4). Asterisks indicate P values smaller than 0.05 by Welch's t-test. Significance levels: * $0.01 \leq P < 0.05$, ** $0.001 \leq P < 0.01$, *** $P < 0.001$.

**Figure S8. OPS antibody mediated complement deposition on Kpn isolates**

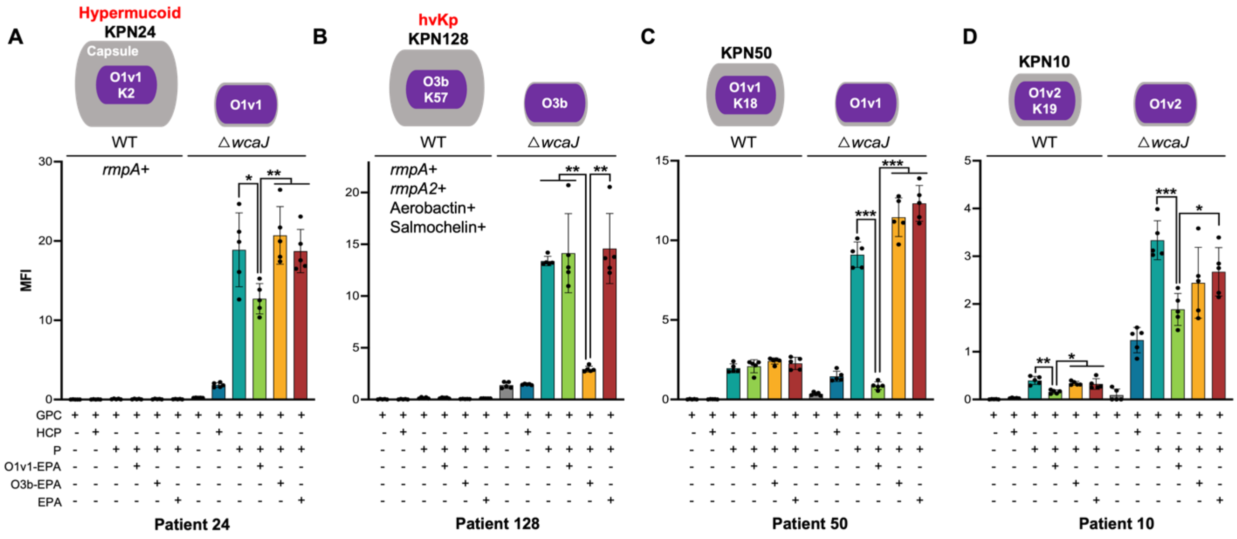


Complement component C3 deposition on Kpn isolates facilitated by OPS antibodies is depicted. Purified immunoglobulins (Ig) from the same plasma samples as described in Figure 4, which include pooled plasma from 36 healthy individuals (HCP) and plasma samples (P) from patients with Kpn BSI were used. After the opsonization process using both untreated and adsorbed Ig (with O1v1-EPA, O3b-EPA, and EPA), guinea pig complement (GPC) was added. The subsequent C3 deposition on both wild-type and capsule-deficient (Δ*wcaJ*) Kpn isolates was measured and presented as median fluorescence intensity (MFI) divided by 1000, on the Y-axis. Asterisks indicate statistical significance with P values smaller than 0.05, determined by Welch's t-test. Significance levels are denoted as follows: * for $0.01 \le P < 0.05$, ** for $0.001 \le P < 0.01$, and *** for $P < 0.001$. The illustration above the figures shows the relative capsule levels in the four Kpn WT strains and their Δ*wcaJ* mutants, based on data presented in Figure S7. It also outlines the OPS and CPS serotypes of Kpn isolates as identified by Kleborate analysis in Table S1, with specific characteristics of KPN24 and the hypervirulent (hvKp) KPN128 depicted in the figures. (N=5)

**Figure S9. MrkA homologues in 69 Kpn isolates**

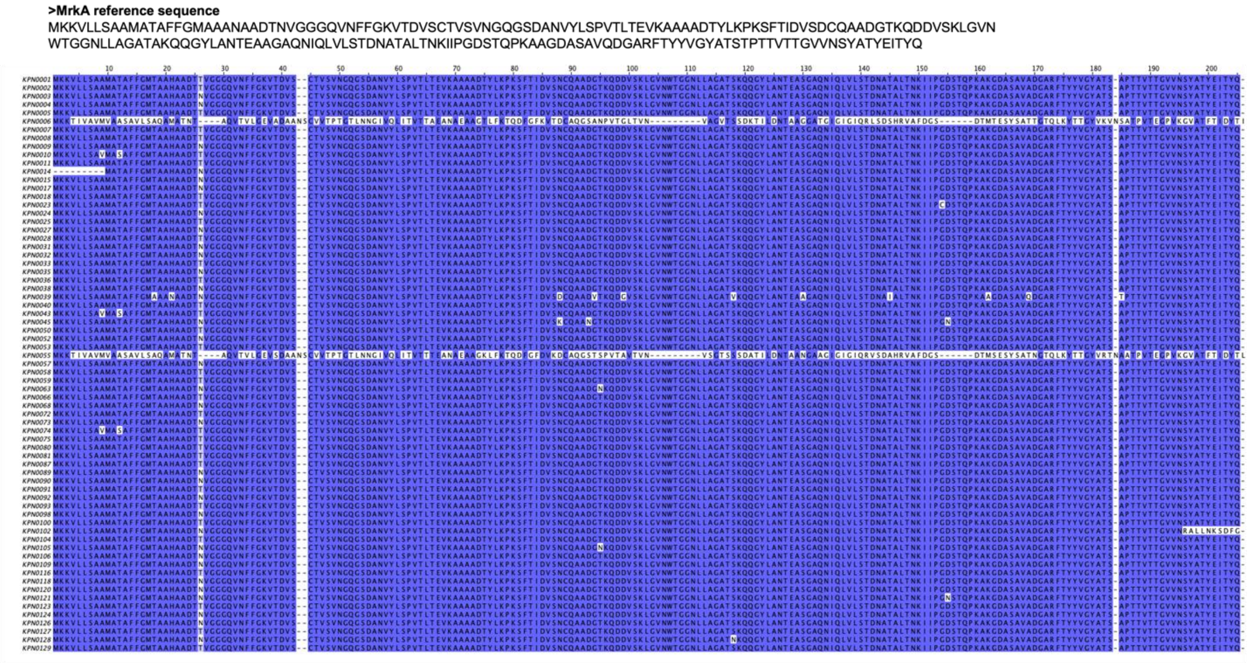


This illustrates the presence of MrkA homologues in 69 Kpn isolates. To identify these homologues, a Blastp analysis was performed using the MrkA reference sequence, in conjunction with the protein sequences obtained from the whole genomes of each Kpn isolate. The criteria for assigning a protein as a representative homologue included a similarity of over 90% and a gene length coverage of more than 85%. The identified MrkA homologues underwent clustering using the CLUSTAL web-based tool, and the resulting clusters were visualized using Jalview. In the visualization, dominant amino acid positions are highlighted in blue, aiding in the identification of conserved residues, while uncolored positions indicate rare occurrences.

**Figure S10. Adsorbed OPS antibody by adding OPS and heat-killed Kpn isolates**

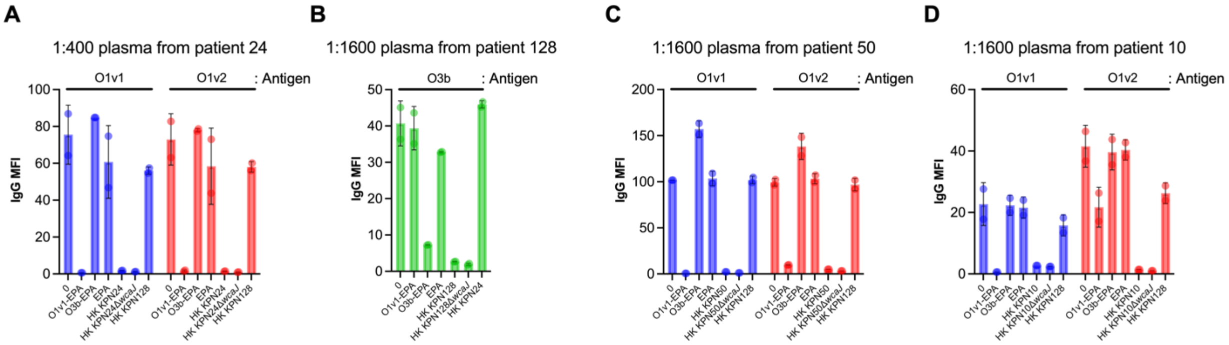


Plasmas from patient 24, 128, 50 and 10 infected with KPN24 (O1v1), KPN128 (O3b), KPN50 (O1v1) and KPN10 (O1v2), at time points with the highest antibody responses to their respective OPSs were used in this assay. These plasmas were diluted to either 1:400 or 1:1600 and incubated with 10 μg/ml of respective antigens (O1v1-EPA, O3b-EPA, and EPA) or 1.5E+6 CFU heat-killed (HK) Kpn isolates or 1.5E+6 CFU HK capsule knockout mutants (Δ*wcaJ*). Both non-adsorbed plasma (denoted as "0") and adsorbed plasmas were utilized in a multiplexed bead assay employing O1v1, O1v2, and O3b antigens. The resulting MFI, divided by 1000, is presented on the Y-axis. Panels **(A)**, **(B)**, **(C)**, and **(D)** correspond to the 1:400 diluted plasma from patient 24 and the 1:1600 diluted plasma from patients 128, 50, and 10, respectively. Negative controls for HK isolates include the use of HK KPN0128 in plasma from patients with O1 Kpn BSI, and HK KPN0024 in plasma from patient with O3b Kpn BSI (*N*=2).

**Table S1. Kleborate results of 69 *Klebsiella pneumoniae* isolates**

| Kpn | Isolation date | K type (CPS) | O type (OPS) ** | *rmpADC* | *rmpA2* | Aerobactin | Salmochelin |
|---|---|---|---|---|---|---|---|
| KPN1 | 7/24/2021 | K14 | O3b | - | - | - | - |
| KPN2 | 7/30/2021 | KL110 | O3b | - | - | - | - |
| KPN3 | 8/24/2021 | K3 | O1v2 | - | - | - | - |
| KPN4 | 9/6/2021 | K38 | O12 | - | - | - | - |
| KPN5 | 9/12/2021 | K14 | O3b | - | - | - | - |
| KPN6 | 9/9/2021 | K31 | O2v1 | - | - | - | - |
| KPN7 | 9/14/2021 | K81 | O1v2 | - | - | - | - |
| KPN8 | 9/18/2021 | K5 | O3b | - | - | - | - |
| KPN9 | 9/18/2021 | K52 | unknown (OL103) | - | - | - | - |
| KPN10 | 9/17/2021 | K19 | O1v2 | - | - | - | - |
| KPN11 | 9/20/2021 | K15 | O4 | - | - | - | - |
| KPN14 | 9/25/2021 | K51 | O12 | - | - | - | - |
| KPN15 | 9/28/2021 | K56 | unknown (OL103) | - | - | - | - |
| KPN17 | 9/27/2021 | KL114 | O1v1 | - | - | - | - |
| KPN18 | 9/29/2021 | K9 | O2v2 | - | - | - | - |
| KPN23 | 10/4/2021 | KL169 | unknown (OL104) | - | - | - | - |
| KPN24 | 10/2/2021 | K2 | O1v1 | *rmp* 3; ICEKp1 | - | - | iro 3 (truncated) |
| KPN25 | 10/6/2021 | K28 | O1v2 | - | - | - | - |
| KPN27 | 10/18/2021 | K21 | O3b | - | - | - | - |
| KPN28 | 10/17/2021 | KL128 | O3b | - | - | - | - |
| KPN31 | 10/24/2021 | K2 | O2v1 -> O1v1 | - | - | - | - |
| KPN32 | 10/27/2021 | KL137 | unknown (OL101) | - | - | - | - |
| KPN33 | 10/28/2021 | KL127 | unknown (OL101) | - | - | - | - |
| KPN35 | 10/29/2021 | K2 | O1v1 | *rmp* 1; KpVP-1 | *rmpA2*_9*-55% | iuc 1 | iro 1 |
| KPN36 | 10/28/2021 | K46 | O3b | - | - | - | - |
| KPN38 | 11/7/2021 | KL103 | O1v1 | - | - | - | - |
| KPN39 | 11/7/2021 | KL110 | O3b | - | - | - | - |
| KPN40 | 11/8/2021 | KL113 | O1v2 | - | - | - | - |
| KPN43 | 11/13/2021 | K62 | O1v2 | - | - | - | - |
| KPN45 | 11/11/2021 | KL174 | O1v2 | - | - | - | - |
| KPN50 | 11/26/2021 | K18 | O1v1 | - | - | - | - |
| KPN52 | 12/1/2021 | K21 | O3b | - | - | - | - |
| KPN53 | 12/3/2021 | K46 | O3b | - | - | - | - |
| KPN55 | 12/6/2021 | K25 | O1v1 | - | - | - | - |
| KPN57 | 12/12/2021 | KL174 | O1v2 | - | - | - | - |
| KPN58 | 12/13/2021 | KL102 | O2v2 | - | - | - | - |

| | | | | | | | |
|---|---|---|---|---|---|---|---|
| KPN59 | 12/16/2021 | KL108 | O1v2 | - | - | - | - |
| KPN63 | 12/31/2021 | K39 | O3/O3a | - | - | - | - |
| KPN66 | 1/6/2022 | KL102 | O2v2 | - | - | - | - |
| KPN68 | 1/16/2022 | KL122 | O2v2 | - | - | - | - |
| KPN72 | 1/19/2022 | K25 | O5 | - | - | - | - |
| KPN73 | 1/19/2022 | K45 | O1v2 | - | - | - | - |
| KPN74 | 1/16/2022 | KL117 | O1v2 | - | - | - | - |
| KPN75 | 1/24/2022 | K14 | O3b | *rmp* 1; KpVP-1 (truncated) | *rmpA2*_5-54% | iuc 1 | - |
| KPN80 | 2/12/2022 | KL102 | O2v2 | - | - | - | - |
| KPN81 | 2/14/2022 | KL145 | O4 | - | - | - | - |
| KPN87 | 3/4/2022 | KL123 | O5 | - | - | - | - |
| KPN89 | 3/17/2022 | K13 | O3b | - | - | - | - |
| KPN90 | 3/26/2022 | KL116 | O2v1 | - | - | - | - |
| KPN91 | 3/26/2022 | K20 | O2v1 | - | - | - | - |
| KPN92 | 3/26/2022 | K13 | O2v1 -> O1v1 | - | - | - | - |
| KPN93 | 3/27/2022 | K11 | O5 | - | - | - | - |
| KPN98 | 5/3/2022 | KL102 | O3b | - | - | - | - |
| KPN100 | 5/4/2022 | K38 | O1v1 | - | - | - | - |
| KPN102 | 5/7/2022 | K17 | O1v1 | - | - | - | - |
| KPN104 | 5/25/2022 | K25 | O5 | - | - | - | - |
| KPN105 | 5/26/2022 | K16 | O1v1 | - | - | - | - |
| KPN106 | 5/29/2022 | K1 | O1v2 | - | - | - | - |
| KPN109 | 6/12/2022 | K52 | unknown (OL101) | - | - | - | - |
| KPN116 | 7/4/2022 | KL102 | O2v2 | - | - | - | - |
| KPN118 | 7/6/2022 | KL155 | unknown (OL101) | - | - | - | - |
| KPN120 | 7/8/2022 | KL151 | O4 | - | - | - | - |
| KPN121 | 7/9/2022 | KL112 | O1v2 | - | - | - | - |
| KPN123 | 7/12/2022 | KL102 | O2v2 | - | - | - | - |
| KPN124 | 7/21/2022 | KL125 | O3b | - | - | - | - |
| KPN126 | 7/23/2022 | K49 | O1v1 | - | - | - | - |
| KPN127 | 7/25/2022 | KL116 | O1v1 | - | - | - | - |
| KPN128 | 7/31/2022 | K57 | O3b | *rmp 1*; KpVP-1 | *rmpA2*_8* | iuc 1 | iro 1 |
| KPN129 | 8/4/2022 | K7 | O3b | *rmp 1*; KpVP-1 | *rmpA2*_8* | iuc 1 | iro 1 |

The table presents data on *Klebsiella pneumoniae* (Kpn) isolates, including their isolation dates, capsular polysaccharide (CPS) and O-specific polysaccharide (OPS) types, as well as the presence or absence of the *rmpA*, *rmpA2*, aerobactin, and salmochelin genes as identified through Kleborate analysis. Isolates classified as hypervirulent Kpn (hvKp), containing either a complete *rmpA* or *rmpA2* gene along with complete aerobactin and salmochelin genes, are highlighted in the table with orange boxes.
** In the case of KPN31 and KPN92, the *wbbY* gene was found across different contigs in the BlastN analysis. Consequently, the O2v1 antigen serotypes of these strains were reclassified as O1v1.

**Table S2. The number of plasma samples used in Figure 2, Figure 3B, and Figure S5**

| IgG | O1v1 | O1v2 | O2v1 | O2v2 | O3/O3a | O3b | O5 | MrkA |
|---|---|---|---|---|---|---|---|---|
| **HC** | 31 | 36 | 36 | 31 | 36 | 31 | 36 | 36 |
| **BC** | 17 | 25 | 25 | 17 | 25 | 17 | 25 | 25 |
| **O1v1** | 10 | 13 | 13 | 10 | 13 | 10 | 13 | 12 |
| **O1v2** | 9 | 13 | 13 | 9 | 13 | 9 | 13 | 13 |
| **O2v1** | 1 | 3 | 3 | 1 | 3 | 1 | 3 | 2 |
| **O2v2** | 4 | 7 | 7 | 4 | 7 | 4 | 7 | 7 |
| **O3/O3a** | 1 | 1 | 1 | 1 | 1 | 1 | 1 | 1 |
| **O3b** | 12 | 16 | 16 | 12 | 16 | 12 | 16 | 16 |
| **O5** | 3 | 4 | 4 | 3 | 4 | 3 | 4 | 4 |
| **other** | 5 | 12 | 12 | 5 | 12 | 5 | 12 | 12 |

| IgM | O1v1 | O1v2 | O2v1 | O2v2 | O3/O3a | O3b | O5 | MrkA |
|---|---|---|---|---|---|---|---|---|
| **HC** | 33 | 36 | 36 | 33 | 36 | 33 | 36 | 36 |
| **BC** | 22 | 25 | 25 | 22 | 25 | 22 | 25 | 25 |
| **O1v1** | 12 | 13 | 13 | 12 | 13 | 12 | 13 | 12 |
| **O1v2** | 10 | 13 | 13 | 10 | 13 | 10 | 13 | 13 |
| **O2v1** | 3 | 3 | 3 | 3 | 3 | 3 | 3 | 2 |
| **O2v2** | 7 | 7 | 7 | 7 | 7 | 7 | 7 | 7 |
| **O3/O3a** | 1 | 1 | 1 | 1 | 1 | 1 | 1 | 1 |
| **O3b** | 15 | 16 | 16 | 15 | 16 | 15 | 16 | 16 |
| **O5** | 4 | 4 | 4 | 4 | 4 | 4 | 4 | 4 |
| **other** | 12 | 12 | 12 | 12 | 12 | 12 | 12 | 12 |

| IgA | O1v1 | O1v2 | O2v1 | O2v2 | O3/O3a | O3b | O5 | MrkA |
|---|---|---|---|---|---|---|---|---|
| **HC** | 33 | 36 | 36 | 33 | 36 | 33 | 36 | 36 |
| **BC** | 18 | 25 | 25 | 18 | 25 | 18 | 25 | 25 |
| **O1v1** | 10 | 13 | 13 | 10 | 13 | 10 | 13 | 12 |
| **O1v2** | 10 | 13 | 13 | 10 | 13 | 10 | 13 | 13 |
| **O2v1** | 2 | 3 | 3 | 2 | 3 | 2 | 3 | 2 |
| **O2v2** | 5 | 7 | 7 | 5 | 7 | 5 | 7 | 7 |
| **O3/O3a** | 1 | 1 | 1 | 1 | 1 | 1 | 1 | 1 |
| **O3b** | 12 | 16 | 16 | 12 | 16 | 12 | 16 | 16 |
| **O5** | 2 | 4 | 4 | 2 | 4 | 2 | 4 | 4 |
| **other** | 7 | 12 | 12 | 7 | 12 | 7 | 12 | 12 |

**Table S3. Primers used in this study**

| | | | |
|---|---|---|---|
| pKPGFP construction | rpsL_pro_F | attcgttaccaatagctatactgatttcgtcag | For amplifying the *rpsL* promoter sequence of pCasKP-apr |
| | rpsL_pro_R | gcctttacgcataaatagctcctggttttag | For amplifying the *rpsL* promoter sequence of pCasKP-apr |
| | sfGFP_F | ccaggagctatttatgcgtaaaggcgaagag | For amplifying the sequence from the start codon of the superfold GFP gene to the *ampR* promoter in pBAD24-sfGFPx1 |
| | sfGFP_R | cgctgatgacatactcttcctttttcaatattattgaag | For amplifying the sequence from the start codon of the superfold GFP gene to the *ampR* promoter in pBAD24-sfGFPx1 |
| | aprR_F | aaaaaggaagagtatgtcatcagcggtggag | For amplifying the *aprR* gene sequence of pCasKP-apr |
| | aprR_R | cttggtctgacagtcagccaatcgactggcg | For amplifying the *aprR* gene sequence of pCasKP-apr |
| | pBadGFP_backbone_F | tcgattggctgactgtcagaccaagtttac | For amplifying the sequence from the end of the *ampR* gene to the end of the *araC* gene in pBAD24-sfGFPx1 |
| | pBadGFP_backbone_R | atcagtatagctattggtaacgaatcagacaattg | For amplifying the sequence from the end of the *ampR* gene to the end of the *araC* gene in pBAD24-sfGFPx1 |
| pDMS197-apr contruction | pDMS197_backbone_F | aataaacaaaacatgagaattgatcctttttgtccggtgttg | For amplifying the plasmid sequence without *tcR* and *tet* promoter sequences in pDMS197 |
| | pDMS197_backbone_R | gattggctgaatggaagccggcggcacc | For amplifying the plasmid sequence without *tcR* and *tet* promoter sequences in pDMS197 |
| | ampRpro_aprR_F | cggcttccattcagccaatcgactggcg | For amplifying the sequence of the *ampR* promoter and *aprR* gene in pKPGFP |
| | ampRpro_aprR_R | attctcatgttttgtttatttttctaaatacattcaaatatgtatccgc | For amplifying the sequence of the *ampR* promoter and *aprR* gene in pKPGFP |
| Primers for confirming inserted sequences in plasmids | pSGKP_spacer_F | ctacgggcctaaagaactaa | For amplifying the sequence near the spacer to confirm the spacer sequence in the pSGKP-spec plasmid |
| | pSGKP_spacer_R | tcacacaggaaacagctatg | For amplifying the sequence near the spacer to confirm the spacer sequence in the pSGKP-spec plasmid |
| | pDMS197-Apr_F | atcaatgattttctggtgcg | For amplifying the sequence near the homologue arm sequences in the pDMS197-apr plasmid |
| | pDMS197-Apr_R | attcaccactccaagaattg | For amplifying the sequence near the homologue arm sequences in the pDMS197-apr plasmid |
| *wcaJ* Deletion in KPN10 | KPN10_wcaJ_spacer_F | tagtgcttatggagacgagatact | KPN10 *wcaJ* spacer |
| | KPN10_wcaJ_spacer_R | aaacagtatctcgtctccataagc | KPN10 *wcaJ* spacer |
| | KPN0010_wcaJ_F | ccgatcttgcaattgaggct | For confirming *wcaJ* deletion in KPN10 genome |
| | KPN0010_wcaJ_R | tcaccagtaacagagtcccc | For confirming *wcaJ* deletion in KPN10 genome |
| | KPN0010_up_F | cgatatcgcatgcggtaccttgctagcccaaatgcagg | Homologue arm sequence, ~0.6 kb upstream of KPN10 *wcaJ* |
| | KPN0010_up_R | tgattgagtgactcccctctctaatcataaaaatg | Homologue arm sequence, ~0.6 kb upstream of KPN10 *wcaJ* |
| | KPN0010_down_F | agaggggagtcactcaatcagtagctgatac | Homologue arm sequence, ~0.6 kb downstream of KPN10 *wcaJ* |
| | KPN0010_down_R | ccttggatcccaagcttctttgataatcaaagcgaccaag | Homologue arm sequence, ~0.6 kb downstream of KPN10 *wcaJ* |
| *wcaJ* Deletion in KPN24 | KPN24_wcaJ_spacer_F | tagtgatatccgcgtgactaaagt | KPN24 *wcaJ* spacer |
| | KPN24_wcaJ_spacer_R | aaacactttagtcacgcggatatc | KPN24 *wcaJ* spacer |
| | KPN0024_wcaJ_F | agaacataagagctacaatatgattttcaa | For confirming *wcaJ* deletion in KPN24 genome |
| | KPN0024_wcaJ_R | ttcggcaatcagctgcat | For confirming *wcaJ* deletion in KPN24 genome |
| | KPN0024_up_F | cgatatcgcatgcggtacctaggatgctaatgtttcgattg | Homologue arm sequence, ~0.6 kb upstream of KPN24 *wcaJ* |
| | KPN0024_up_R | gagcatctagaattcaatcactcatttataaacaag | Homologue arm sequence, ~0.6 kb upstream of KPN24 *wcaJ* |
| | KPN0024_down_F | tgattgaattctagatgctccttaagacaag | Homologue arm sequence, ~0.6 kb downstream of KPN24 *wcaJ* |
| | KPN0024_down_R | ccttggatcccaagcttctttataggtcacacacggctc | Homologue arm sequence, ~0.6 kb downstream of KPN24 *wcaJ* |
| *wcaJ* Deletion in KPN50 | KPN50_wcaJ_spacer_F | tagtgtgcctaaagtttcgatcca | KPN50 *wcaJ* spacer |
| | KPN50_wcaJ_spacer_R | aaactggatcgaaactttaggcac | KPN50 *wcaJ* spacer |

| | | | |
|---|---|---|---|
| | KPN0050_wcaJ_F | tgatagtcgatacgcctcct | For confirming *wcaJ* deletion in KPN50 genome |
| | KPN0050_wcaJ_R | tgggctatagggtttgcatc | For confirming *wcaJ* deletion in KPN50 genome |
| | KPN0050_wcaJ_dsDNA | acatcaattactgtgattaataaaatatataagtaagagggatatcttttttaaaaatctcacagtgagttttcaacgaaatgaaatgt | 90 bp of *wcaJ* upstream 45 bp and *wcaJ* downstream 45 bp of KPN50 |
| *wcaJ* Deletion in KPN128 | KPN0128_up_F | cgatatcgcatgcggtacctgagctagtcctaatgcgg | Homologue arm sequence, ~0.6 kb upstream of KPN128 *wcaJ* |
| | KPN0128_up_R | gcataatgaaagttacacgcctacttatg | Homologue arm sequence, ~0.6 kb upstream of KPN128 *wcaJ* |
| | KPN0128_down_F | gcgtgtaactttcattatgcttggataaaaacaatg | Homologue arm sequence, ~0.6 kb downstream of KPN128 *wcaJ* |
| | KPN0128_down_R | ccttggatcccaagcttcttcccgtgccattgaaaactc | Homologue arm sequence, ~0.6 kb downstream of KPN128 *wcaJ* |
| | KPN0128_wcaJ_F | ttgaaaacccagcagatttg | For confirming *wcaJ* deletion in KPN128 genome |
| | KPN0128_wcaJ_R | agcgtatgagttgtctgaaatca | For confirming *wcaJ* deletion in KPN128 genome |
